# DisCell: Separating Spillover from Niche Response in Spatial Transcriptomics

**DOI:** 10.64898/2026.11.04.764463

**Authors:** Ricky Molén, Hosein Toosi, Pedro F. Ferreira, Jens Lagergren

## Abstract

Spatial transcriptomics at single-cell resolution measures every cell in its place in the tissue, but diffusion and segmentation errors also add transcripts of neighboring cells to a cell’s counts. This spillover makes a cell resemble its neighbors, and a response to the niche makes cells in the same niche resemble each other. Both tie a cell’s expression to its surroundings. The count data alone cannot tell the two apart.

Findings from single-cell transcriptomes, such as a gene program that is higher in one niche than in another or signaling genes that change depending on the neighborhood, may therefore be artifacts of spillover. Current models describe a cell by its intrinsic state together with either its niche response or spillover, but none models all three.

We present DisCell, the first method that models all three components jointly and measures how robust each finding is to spillover. DisCell is a generative model that describes single-cell gene expression profiles in terms of two latent variables, an intrinsic state and a response to the niche, and adds spillover from its neighbors as a separate term. DisCell fits the model at increasing levels of spillover and recomputes each finding on the spillover-free expression to see how much spillover it withstands. DisCell also estimates the most spillover a section can contain, and a finding that holds even at that amount is not explained by spillover.

In simulated tissue, DisCell recovers each cell’s intrinsic state and response and tells findings created by spillover apart from genuine responses. On five sections of ovarian and lung tissue, the intrinsic state keeps more of each cell’s own properties, measured by its cell-cycle phase, than any competing method that holds as little of its neighborhood. Within each cell type, the model partitions the variation of each gene into intrinsic state, niche response and spillover. Using DisCell, we rank findings about niches by their robustness to spillover and confirm that a previously reported finding withstands spillover, that esophageal cancer cells in niches infiltrated by immune or stromal cells express more of an epithelial–mesenchymal transition program. DisCell also predicts what the gene expression of a cell type would be in another niche, with its spillover part separated, at lower error on most tasks than the competing method built for this purpose. Training the model takes under ten minutes on one consumer GPU for sections of up to 400,000 cells.

## 1 Introduction

The impact of a cell’s surroundings on its state is a main question that spatial transcriptomics at single-cell resolution is uniquely poised to answer. Imaging-based platforms detect individual transcripts of thousands of genes in a tissue section and assign each to a cell, so every cell gets a count per gene [1, 2]. These counts come from three sources. The first is the cell’s *intrinsic state*, which comprises its type, its cell-cycle phase and any other property that does not depend on its surroundings. The second is the cell’s *niche response*, carried by pathways that sense the surroundings and change transcription, so it varies with the neighborhood. The third is *spillover*, a measurement artifact in which transcripts of neighboring cells are assigned to the cell, because segmentation of densely packed tissue is imperfect and because transcripts can diffuse away from the cell that made them before they are detected [3]. Spillover is common: counting the transcripts of other cell types’ marker genes, Yang et al. [4] found more than 5% of counts in such genes in 3 to 10% of cells measured in single-cell RNA sequencing, but in 47 to 73% of immune cells and fibroblasts on a Xenium section of lung cancer.

A cell that receives its neighbors’ transcripts acquires their markers, and a cell that responds to its neighbors shifts its programs, so the response and spillover are confounded. Spillover in a cell grows with the local abundance of the neighboring type [5], and models of niche effects describe the response as a function of the same neighbor composition [6]. A shift in genes that the neighbors do not express cannot be spillover, however strong the spillover is. A shift toward the neighbors’ profile is different; it can be a response to the neighbors or spillover from them, and since a flexible model fits the counts equally well either way, how much of the shift is called response and how much spillover is decided by the modeling assumptions, not by the data.

Spillover can alone produce apparent cell–cell interaction signals [7], so any inferred niche program or interaction between neighboring cells [8, 9] may be an artifact of spillover. However, separating biological signal from segmentation artifacts in patterns between neighboring cells remains an open problem [10]. Whether such a result is real therefore depends on how much of the shift toward the neighbors is attributed to spillover. Correcting the counts before an analysis does not remove this dependence, because each correction removes one assumed amount of spillover and the counts alone cannot check whether that amount of spillover is right. Corrections that estimate the amount, such as DeLeakage [3], do so by assuming that all cells of a specific type share one expression profile, which leaves no room for a niche response.

### Related work

Per-gene models split a gene’s variance into spatial and non-spatial parts [11, 12] or regress each gene on niche covariates [13, 14], and cell-level models describe a cell’s expression by latent factors that mix intrinsic and spatial influences [15] or split it into an intrinsic and a neighbor-dependent part [16, 17]. Contamination is removed separately, before such analyses [7, 18–24]. Among deep generative models of the whole cell, each models either the niche response or spillover, but not both (Table 1). resolVI [25] models spillover but has no niche term, so a response that shifts a cell toward its neighbors can only be assigned to the cell’s own state or to spillover. SIMVI [26], MintFlow [27] and Cellina [28] split a cell’s expression into intrinsic and niche-dependent factors but do not model spillover, and MintFlow and Cellina list segmentation errors, a cause of spillover, as a limitation. Other models of niche effects ignore spillover, filter it after the fit or test it at fixed levels [6, 29–32].

**Table 1:**
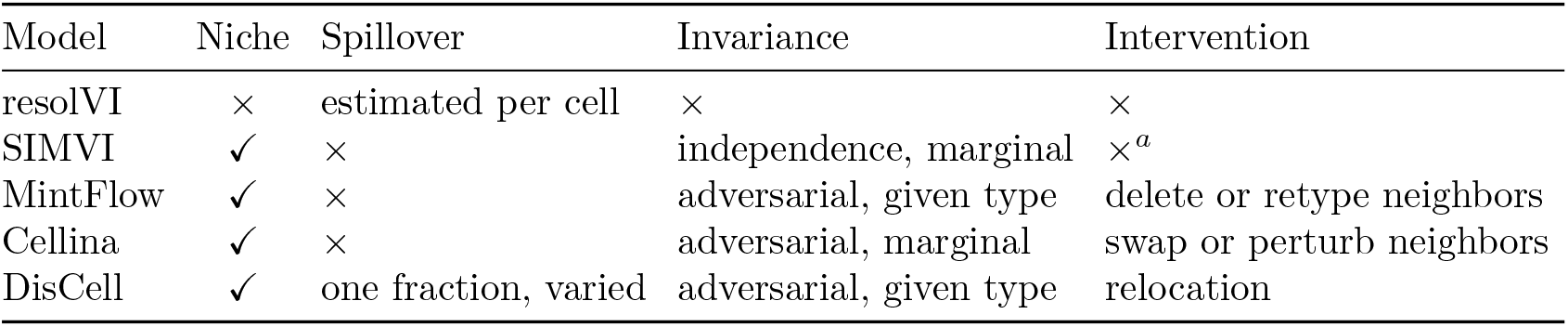
The generative models most similar to DisCell. The columns give whether misassigned transcripts are part of the model (Spillover), how the intrinsic latent is kept free of the niche (Invariance), and which change of the neighborhood the model can simulate (Intervention). MintFlow samples new cells for the edited tissue, while Cellina and DisCell predict the observed cells in a new neighborhood. Cellina can also change genes in a cell’s neighbors, and DisCell splits the predicted shift into response and spillover. ^*a*^SIMVI instead estimates a spatial treatment effect.

| Model | Niche | Spillover | Invariance | Intervention |
|---|---|---|---|---|
| resolVI | × | estimated per cell | × | × |
| SIMVI | ✓ | × | independence, marginal | ×<sup>*a*</sup> |
| MintFlow | ✓ | × | adversarial, given type | delete or retype neighbors |
| Cellina | ✓ | × | adversarial, marginal | swap or perturb neighbors |
| DisCell | ✓ | one fraction, varied | adversarial, given type | relocation |

### Our approach

Here, we present DisCell, a generative model that describes each cell by an intrinsic state and a response to its niche, and mixes into its counts a fraction *κ* of its neighbors’ expression as spillover. Because the counts bound *κ* from above but never from below (Proposition 1), DisCell does not commit to one amount of spillover. It is refitted over a range of *κ*, and at each value it splits every cell’s expected expression into what is the cell’s own and what spills in from its neighbors, so any finding that is computed from the cells’ expression can be recomputed on the spillover-free expression. We call such a result a *finding*, for example a gene program that is higher in one niche than in another, or signaling genes that change with a cell’s neighbors. The smallest amount of spillover at which a finding no longer holds is its *breakdown point*. Spillover only adds the neighbors’ transcripts, so removing too much of it would take away more of some gene than a cell type has. With DisCell we estimate, for each section, the largest spillover fraction that avoids this, so a user can see which findings still hold when that much spillover is removed. DisCell shares components with earlier models. Its spillover term is a mixture like those that remove contamination [19, 25], and its adversary is like those of MintFlow and of perturbation models [27, 33, 34]. DisCell models both niche response and spillover jointly and uses this to present a test that every finding must pass.

### Contributions

- **DisCell separates a cell’s intrinsic state from its niche response and spillover**. Its intrinsic state keeps cell identity and the cell cycle, and no competing method keeps more of the cycle while holding less of the neighborhood. Within a cell type it partitions the variation of each gene into intrinsic, niche response and spillover (Sec. 3.1).
- **Every finding computed from the cells’ expression gets a breakdown point, read against an estimated bound on spillover**. This ranks findings by robustness and says which ones spillover of the modeled kind cannot explain, including a published finding in esophageal cancer (Sec. 3.2).
- **DisCell predicts how a cell type would change in another niche, and how much of that change is spillover**. For cells withheld from every model’s training, its predicted shifts have lower error than each tested form of Cellina on most tasks (Sec. 3.3).

DisCell needs only counts, cell positions and cell types. Code is available at https://github.com/molen89/DisCell.

## 2 Methods

### 2.1 The model

DisCell is fitted to one section at a time. For each segmented cell *i* we observe its counts **x**_*i*_ over *G* genes, with total *ℓ*_*i*_, its cell type *t*_*i*_, its position and the morphology image around it. Two cells are neighbors, *j* ∈ *N* _*i*_, when they share a face of the cells’ Voronoi tessellation (S4). Three quantities describe the surroundings without using the cell’s own counts. The composition **y**_*i*_ is the share of each cell type among the neighbors. The image context **Φ**_*i*_ embeds the morphology image around the cell, with the cell masked (Fig. 1b). Any image encoder can produce it, and we use the foundation model KRONOS [35]. The niche descriptor **c**_*i*_ joins the image context to an attention layer over the neighbors’ types, queried by the cell’s type. DisCell describes the *i*th cell by two latent variables, its intrinsic state **z**_*i*_ and its niche response **w**_*i*_ (Fig. 1c),

**Fig. 1:**
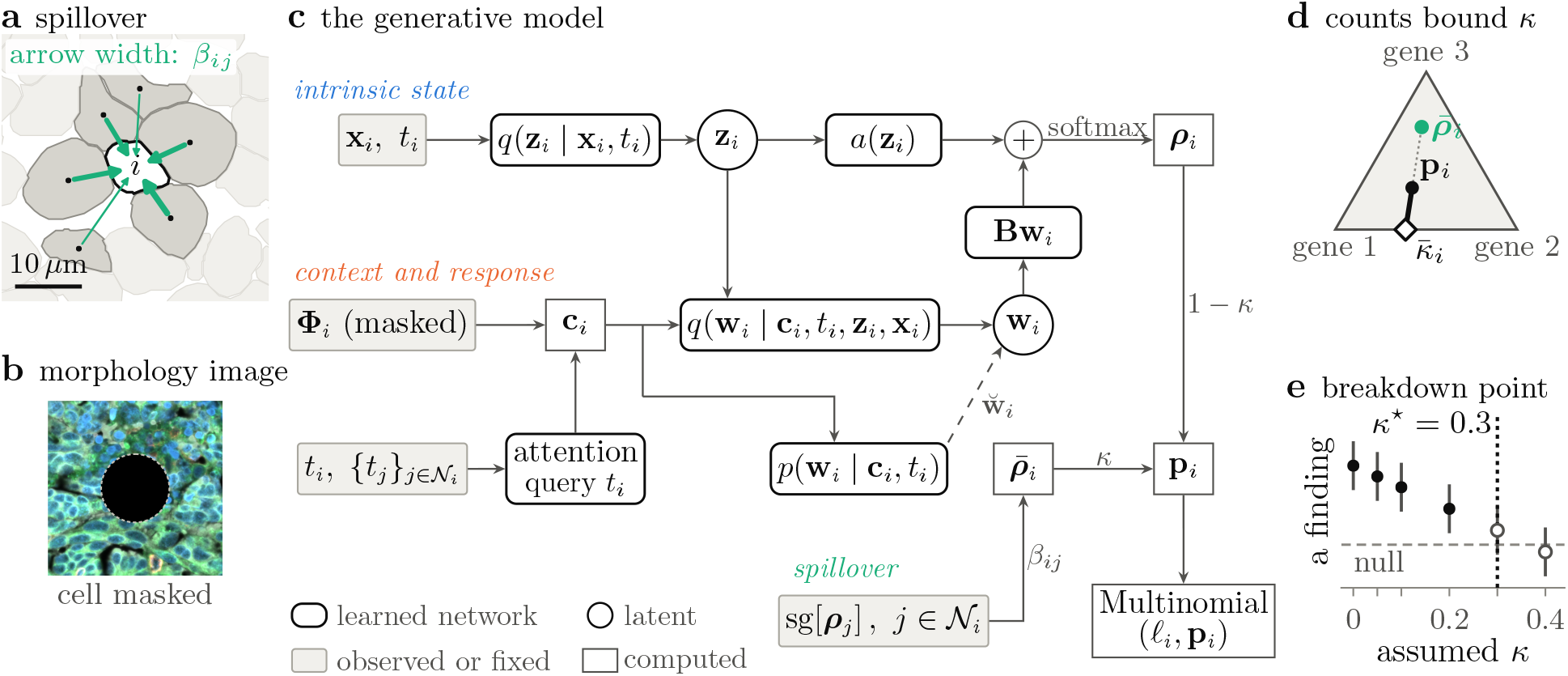
DisCell overview. **a**, Spillover on segmented cells of an ovarian cancer section. **b**, The morphology image around a cell, with the cell masked, used to create the image context **Φ**_*i*_. **c**, The model and how it is fitted (Secs. 2.1 and 2.2). The intrinsic latent state **z**_*i*_, modeled as a Gaussian with mean ***µ***_*z,i*_, is encoded from the cell’s counts and type and the response **w**_*i*_ from its niche descriptor **c**_*i*_ as well as the inputs and the output of the encoder for **z**_*i*_. Both **z**_*i*_ and **w**_*i*_ are used to decode the cell’s own profile ***ρ***_*i*_, which is mixed with its neighbors’ profile 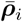 at the fraction *κ*. Here sg is a stop-gradient, *p*(**w**_*i*_ | **c**_*i*_, *t*_*i*_) the prior of the response, 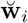 a response drawn from it, **B** the matrix of gene programs and *ℓ*_*i*_ the cell’s total count. **d**, On the simplex of three genes, each assumed *κ* gives a clean profile on the solid segment that reproduces the cell’s measured profile **p**_*i*_ exactly. The segment ends at 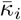 (diamond), where one gene’s share reaches zero, so the counts bound *κ* only from above (Proposition 1). **e**, A finding refitted at each assumed *κ*, with its breakdown point *κ*^*⋆*^ (Definition 1). Filled points hold, open points no longer hold, and the dashed line is the null.

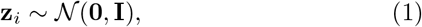

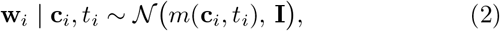

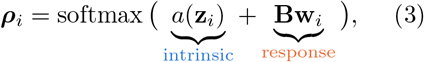

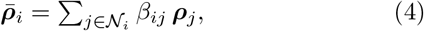

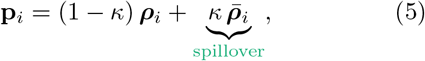

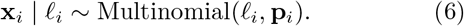

The intrinsic state **z**_*i*_ holds the cell’s own properties, which it would keep in any niche, and a network *a* maps it to log-shares over genes. The response **w**_*i*_ is how far the cell’s expression has shifted along the gene programs in the columns of **B**, and a network *m* gives its prior mean *m*(**c**_*i*_, *t*_*i*_) as the response expected of a cell of type *t*_*i*_ in niche **c**_*i*_. Together, *a* and **B** form the decoder, which gives the cell’s own, *clean* expression profile ***ρ***_*i*_, its expected share of each gene’s transcripts without spillover. In the spillover term, the neighbors’ clean profiles are averaged into 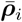 with weights 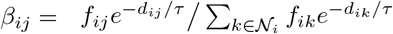, where *f*_*ij*_ is the length of the face two cells share, *d*_*ij*_ the distance between their centers and *τ* a decay length. The fraction *κ* is therefore the share of a cell’s expected counts that come from its neighbors. It is the same for every gene and cell of a section and fixed for a fit, and DisCell is refitted over a range of *κ* values (Sec. 2.4).

### 2.2 Fitting the model

Each latent has its own encoder, a network that gives the cell a diagonal Gaussian posterior 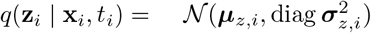 for the intrinsic state and 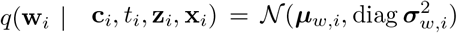 for the response.

The encoder of the intrinsic state reads only the cell’s own counts and type, never its niche. Over batches *S* of cells, DisCell maximizes

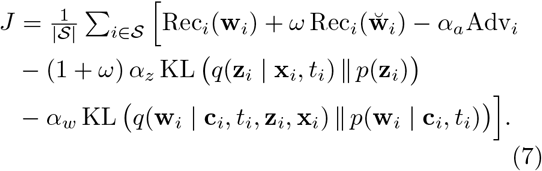

Here Rec_*i*_(**w**) scores how well the cell’s counts are reconstructed when its response is **w**. It is their log-likelihood per transcript, 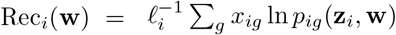, with *p*_*ig*_(**z**_*i*_, **w**) gene *g*’s share in **p**_*i*_ (Eq. (5)) computed with that response. The two Kullback–Leibler divergences (KL), weighted by *α*_*z*_ and *α*_*w*_, pull the posteriors toward their priors, N (**0, I**) and N (*m*(**c**_*i*_, *t*_*i*_), **I**). *J* is a weighted evidence lower bound (S2.1) with two additions. The first, 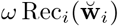, reconstructs the cell a second time with a response 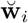 drawn from the prior (the dashed arrow in Fig. 1c), adapting the two-path objective of Slavutsky et al. [36]. The intrinsic state, rather than the response, must then carry what sets the cell apart from others of its type in its niche, and its divergence is counted once per reconstruction, hence the factor 1 + *ω*. The second addition is a penalty, with weight *α*_*a*_, on the adversary’s advantage Adv_*i*_, which the next paragraph defines in Eq. (9).

#### An adversary for the niche

Within each cell type, the intrinsic state should carry no information about the niche. For a cell drawn at random from the section, with intrinsic state **z**, composition **y**, image context **Φ** and type *t*, this means

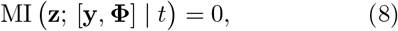

where MI(·;·|*t*) is the mutual information conditional on the type. It is the average of the mutual information within each type, so it is zero only if it is zero within every type. The condition is given the type because cell types cluster in space, and an intrinsic state blind to the niche overall would have to lose part of the type [37]. The adversary is two small networks that read only the posterior mean ***µ***_*z,i*_ and the type. From these they predict the composition as **ŷ**_*i*_ and the image niche as **ê**_*i*_, where **e**_*i*_ softly assigns **Φ**_*i*_ to clusters. Its advantage is how much better they do than the type’s averages 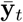 and **ē**_*t*_, with *λ*_*y*_ weighting the composition,

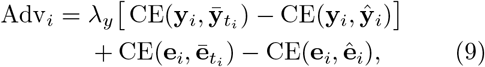

with CE(**a, b**) = −Σ_*k*_ *a*_*k*_ ln *b*_*k*_ the cross-entropy. Averaged over cells, each of its two terms is a lower bound on the information ***µ***_*z,i*_ adds about the niche beyond the type, so it is zero when the adversary finds none, as Eq. (8) requires. The rest of the model maximizes *J*, while the adversary’s networks maximize Adv_*i*_ with ***µ***_*z,i*_ held fixed. The two take alternating gradient steps on each batch, with separate optimizers, as in Cellina [28] and similar to MintFlow [27].

That the adversary can no longer predict the niche does not show that the information is gone, since another predictor may still find it [38], so after training a probe, fitted on training tiles, checks the result on held-out tiles. It reports the share of the neighborhood’s variation within types that it recovers from **z**_*i*_ beyond chance (S3.1), for every method in Fig. 2.

**Fig. 2:**
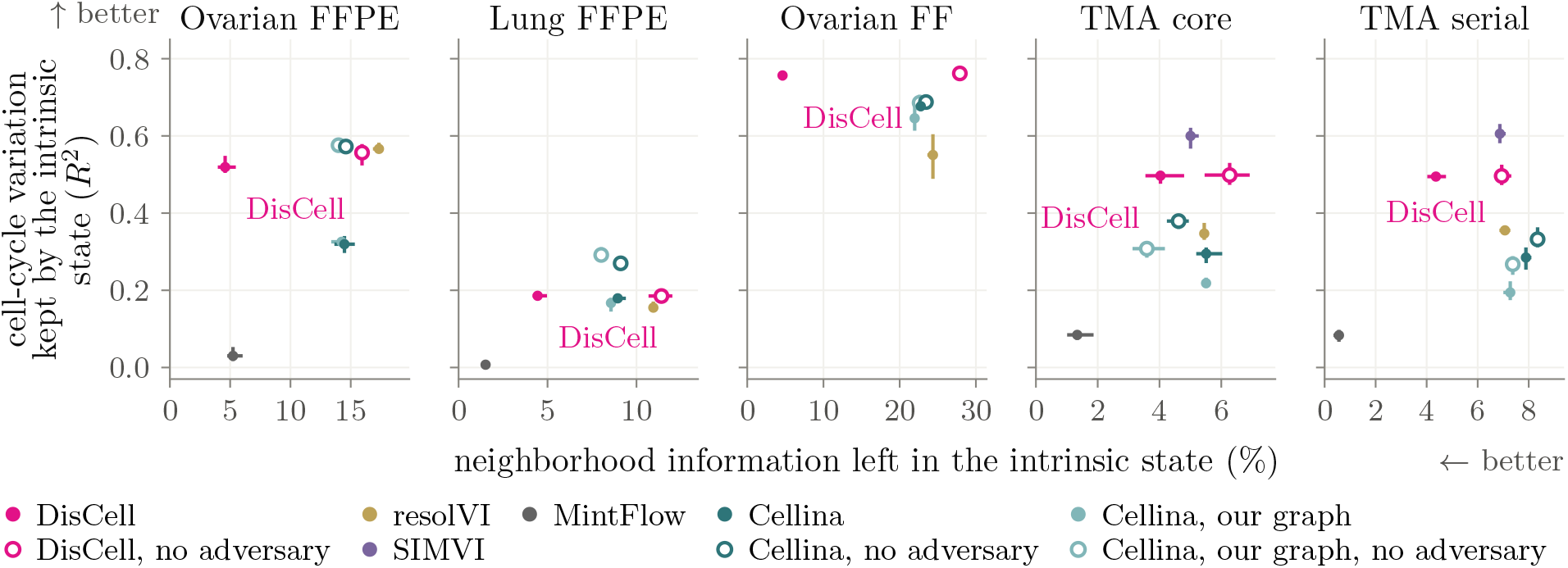
Neighborhood information and cell-cycle variation in the intrinsic state of each method, on held-out cells. For the other methods, the intrinsic state is the corresponding latent variable. Horizontal: within-type variation of neighbor composition recovered by a probe, beyond chance; vertical: within-type cell-cycle *R*^2^ (S3.1). Means over three seeds, with lines spanning the range. On the serial section, resolVI and SIMVI were fitted on that section, and the other methods were evaluated with the models fitted to the TMA core. Missing methods could not be run on that section (S4.1).

### 2.3 What the counts can tell about spillover

The counts bound *κ* only from above, since removing too much of the neighbors’ profile would leave some gene with a negative share (Fig. 1d).

#### Proposition 1.

*For a cell with at least one neighbor, hold the neighbors’ clean profiles, and hence*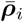, *fixed, and let* 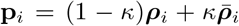 *with* ***ρ***_*i*_ *in the simplex. A fraction κ*^*′*^ ∈ [0, 1) *reproduces* **p**_*i*_ *with a clean profile in the simplex if and only if* 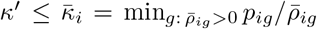. *The true κ satisfies*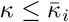.

For one cell the bound is too noisy to use, so we pool it over the cells of a type, with a confidence limit so that a gene with few counts cannot set it by chance. The counts-only estimate assumes the neighbors’ observed profiles are free of spillover and needs no model, and the model-based estimate uses the clean profiles of a fitted model (S2.2). A section’s 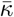 is the largest value over its cell types, the loosest of the type bounds, so it rules out as little spillover as the data permit. 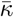 bounds spillover of the modeled kind only. It is not a measurement of the physical share of misassigned transcripts.

### 2.4 The breakdown point

We fit the model independently at each *κ* ∈ {0, 0.05, 0.1, 0.2, 0.3, 0.4 }, three times with different random seeds and every other setting fixed. This is a sensitivity analysis with *κ* as the sensitivity parameter, in the tradition of tipping-point and E-value analyses of unmeasured confounding [39, 40].

#### Definition 1

(Breakdown point). State a finding as a number *r* that is zero in expectation when the effect is absent, positive in the claimed direction and negative in the opposite one. At each *κ*, let CI_*κ*_(*r*) be a confidence interval for the mean of *r* over the three fits. The finding holds at *κ* if CI_*κ*_(*r*) lies entirely above zero. Its breakdown point *κ*^*⋆*^ is the smallest tested *κ* at which it does not hold, and is reported as above the largest tested value if it holds at all of them. A finding that does not hold at *κ* = 0 has no *κ*^*⋆*^.

The interval is a bootstrap over both sources of variation. Each draw picks three of the fits with replacement, resamples 200 *µ*m squares of tissue within each, and averages *r*. Its level is corrected for the number of findings tested on the section (S3.4). *κ*^*⋆*^ neither estimates *κ* nor certifies a finding. It says how much spillover of the modeled kind must be assumed to lose the finding. Read against the section’s bound 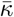, it gives one of three outcomes (Fig. 5). If a finding still holds at a tested *κ* at or above 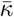, such spillover within the bound does not explain it. If it breaks before holding at any tested *κ* at or above 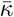, it may be an artifact of spillover. If it holds at every tested *κ* but 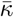 lies above the largest, it is undecided, since it could still break between the two.

### 2.5 Data, comparisons and use

We fit DisCell to four Xenium [2] sections with a panel of about five thousand genes. Three are whole-slide sections, of ovarian cancer fixed in formalin and embedded in paraffin (FFPE) [41], lung cancer FFPE [42] and fresh-frozen (FF) ovarian cancer [43], and the fourth is one core of a pediatric lung tissue microarray (TMA) [44], with 69,000 to 1.16 million cells per section. A serial section of the TMA core, on a second slide, is never fitted and is evaluated only with the models fitted to the core. Only Ovarian FFPE came with curated cell types, so on the other sections we named clusters by their marker genes, and on the TMA also by correlation with lung reference atlases (S4). Each cell carries a lineage label, and finer states and locations are merged into their lineage. Two sections from an independent esophageal cancer study [45] re-test a published finding (S5.3). Their public release has no images. 15% of each section’s spatial tiles are held out. Results at a single *κ* use *κ* = 0.1, a moderate value on the scale marker counts suggest [4].

We compare with resolVI, SIMVI, MintFlow and Cellina at the model settings of their papers, scoring every method’s intrinsic latent on the same held-out cells (S4.1). They were fitted on all cells, including our held-out ones, which favors them. In the relocation test of Fig. 7, no model saw the withheld cells.

## 3 Results

The results cover four Xenium sections (Ovarian FFPE, Lung FFPE, Ovarian FF and a TMA core) and two published esophageal cancer sections (Sec. 2.5). We first test whether the intrinsic state and the response separate as intended (Sec. 3.1), then use breakdown points to assess findings about niches (Sec. 3.2), and finally test relocation on cells withheld from training (Sec. 3.3).

### 3.1 DisCell separates the intrinsic state from the niche

#### The intrinsic state contains cell-cycle variation and little neighborhood information

We measured on held-out cells how much neighborhood information and how much within-type cell-cycle variation the intrinsic state contains (S3.1). The cell-cycle measure is included to exclude intrinsic states that are free of the niche only because they are uninformative. We quantified neighborhood information with a probe fitted to predict the composition **y**_*i*_ from ***µ***_*z,i*_ and *t*_*i*_, as in the adversary, and cell-cycle variation as the *R*^2^ of a ridge regression of the S and G2/M scores [46], centered per type, on ***µ***_*z,i*_, in the tenth of held-out cells with the highest scores. We computed both measures for DisCell and for the intrinsic latents of resolVI, SIMVI, MintFlow and Cellina (Fig. 2, Table S4). No intrinsic latent contains both more cell-cycle variation and less neighborhood information than that of DisCell on any section, including the serial section. The intrinsic latent of resolVI, which models spillover but no niche, contains two to five times as much neighborhood information as that of DisCell on the whole-slide sections, that of MintFlow contains less on Lung FFPE and the TMA sections but almost no cell-cycle variation, and that of Cellina contains more in every form and section except one. The results on the serial section, which was not used in training, are similar to those on the core. On sections simulated from the model, the intrinsic state recovers the true cell types and the response the true response, and without the adversary the intrinsic state contains the simulated niche effect (S5.1).

#### Within each cell type, DisCell partitions each gene’s variation into intrinsic state, niche response and spillover

We then decomposed the variation of each gene across the cells of each type into contributions from the intrinsic state, the response and spillover, and assigned the gene to the dominant contribution (S3.5). We show each gene as a point in a triangle whose corners correspond to the three contributions (Fig. 3). Most genes of Ovarian FFPE lie near the intrinsic corner, and the remaining genes lie toward the response or toward spillover. Because the contributions are computed over all cells of a type, a gene with a response or spillover confined to a few of these cells can still lie near the intrinsic corner, and we show in Fig. 4 where in the tissue each contribution is large. *CTHRC1*, a gene of cancer-associated fibroblasts, is assigned to the niche response in fibroblasts, genes specific to fibroblasts such as *DCN* to the intrinsic state, and the endothelial tip-cell gene *ESM1* to spillover. Macrophage markers such as *CD163* and *MS4A6A* are assigned to spillover in pericytes, as are smooth-muscle genes such as *MYH11* and *CNN1* in macrophages and endothelial cells. The genes assigned to spillover are thus markers of neighboring cell types, as expected, and every assignment to the response or to spillover in these four types holds beyond the section’s estimated bound on spillover.

**Fig. 3:**
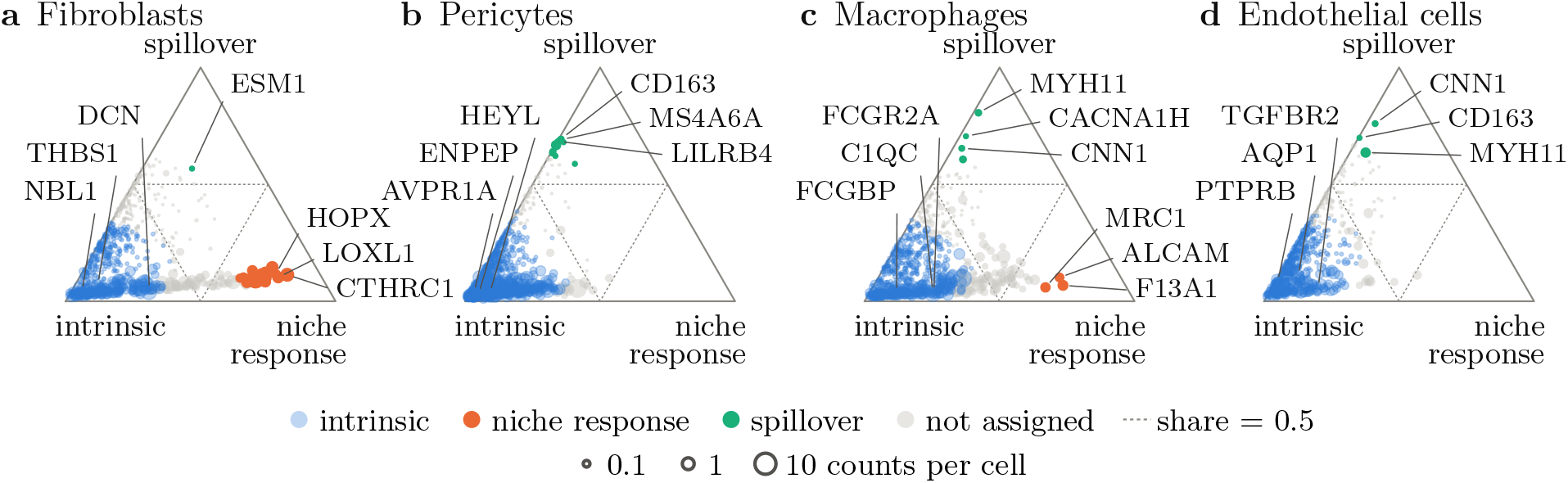
Decomposition of the within-type variation of each gene into intrinsic, niche-response and spillover contributions, on Ovarian FFPE at. *κ* = 0.1. Each point is a gene, positioned by the three shares (means over three seeds, S3.5), so a gene near a corner is mostly explained by that contribution. Genes are colored when one share exceeds one half and is at least twice each of the other two, and gray otherwise. Every assignment to the response or to spillover shown holds beyond the estimated bound 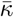 of the section. The shares are computed on the expected expression of the model, which excludes the counting noise that makes up most of the variation of the observed counts.

**Fig. 4:**
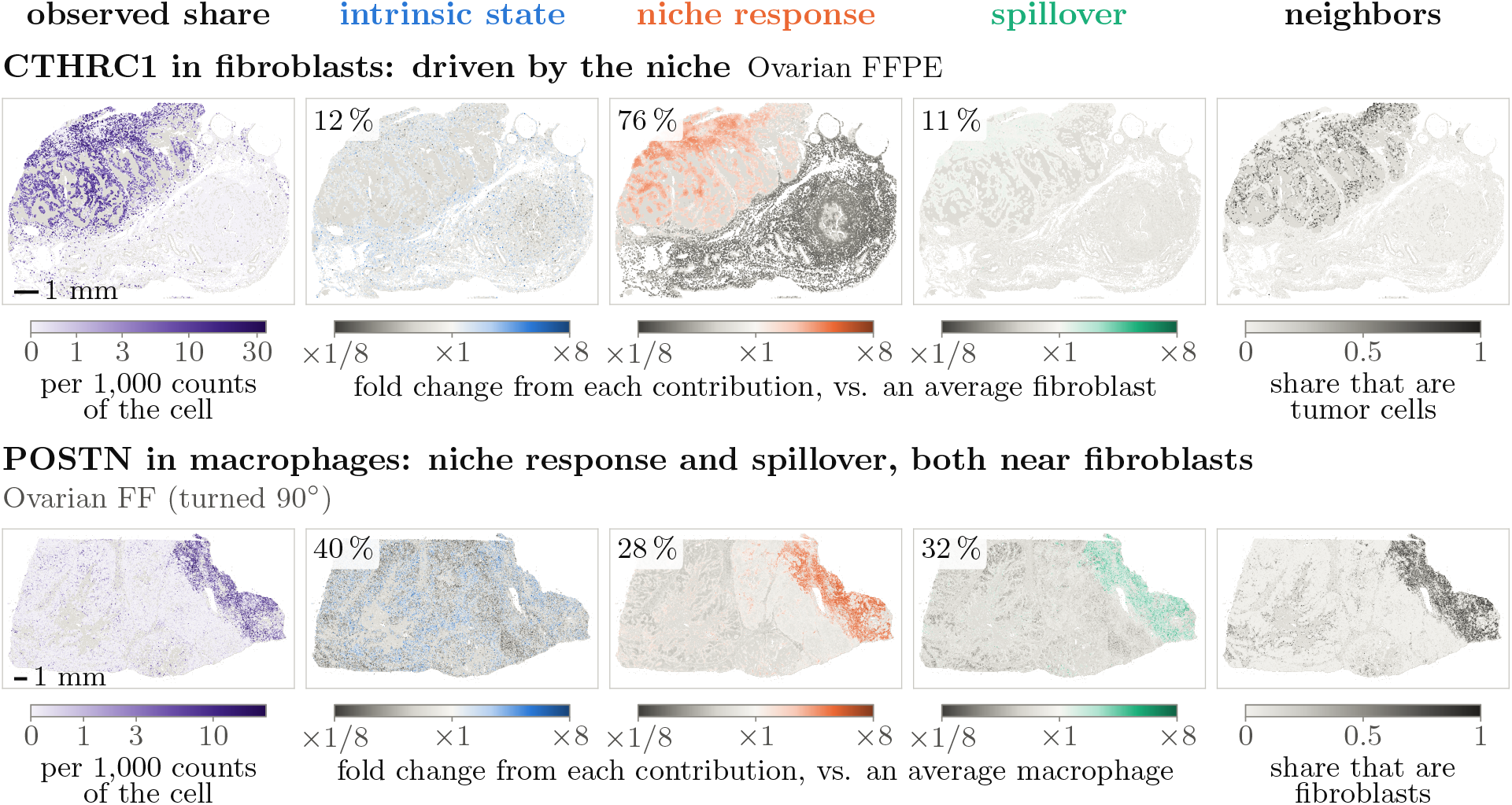
Intrinsic, niche-response and spillover contributions to two genes over whole sections of ovarian cancer. Each row shows one gene in one cell type; dots are cells of that type at the cell positions, and light gray dots are all other cells. First column: observed share of the gene. Columns two to four: fold change of the gene due to the intrinsic state, the niche response and spillover, relative to an average cell of the type, either increased (color) or decreased (gray); percentages are the shares of the within-type variation of the gene, as in Fig. 3, averaged over three seeds. Last column: share of neighbors of the named type, weighted as in the spillover term. Both rows use *κ* = 0.1, and the maps show one seed.

#### The decomposition of gene variation agrees with known biology

Then we mapped the intrinsic, response and spillover contributions over the tissue (Fig. 4). The niche-response contribution to *CTHRC1* in fibroblasts is positive near the tumor and negative in the stroma, following the share of tumor neighbors, and accounts for 76% of the variation of *CTHRC1* in fibroblasts, as expected for a marker of fibroblasts at the leading edge of tumors [47]. *POSTN*, a marker of matrix-remodeling cancer-associated fibroblasts [48], is detected in macrophages of Ovarian FF where fibroblasts are dense. The variation of *POSTN* in macrophages is split between spillover from those fibroblasts (32%) and a niche response (28%) at *κ* = 0.1, but spillover accounts for 76% at *κ* = 0.4, which is below the estimated bound of 0.49 for this section.

### 3.2 Breakdown points flag the findings that spillover could explain

#### In simulation, the breakdown point separates spillover from response

The true cell types and responses of sections simulated with *κ* = 0.2 are recovered about equally well at any assumed *κ* from 0 to 0.4 (S5.1), as expected since *κ* is bounded only from above by the counts. We therefore tested whether the breakdown point separates findings without knowledge of *κ*, on simulated sections of 30,000 cells with two known findings at a true *κ* of 0, 0.1 or 0.2: a difference between niches created only by spillover, and a genuine response. The spillover-only difference does not hold even at *κ* = 0 without simulated spillover, and breaks at *κ* = 0.05 and 0.1, before the true *κ*, with spillover at 0.1 and 0.2. The genuine response holds beyond 0.4 in every simulation.

#### Most findings on tissue hold beyond the estimated bound

We then tested three findings about niche responses (Fig. 5, defined in Table S1). One comes from relocation (Sec. 3.3): the response part of a predicted shift points in the direction of the observed shift once its spillover part is removed. Two are criteria from earlier work: ligand and receptor genes take a larger share of the response than other genes (the signaling-gene lean) [27], and the response changes with distance to the tumor (the tumor axis) [26]. All three findings hold beyond *κ* = 0.4 on Ovarian FFPE, whose bound is about 0.18, so spillover of the modeled kind cannot explain any of them. The two findings tested on the TMA core and its serial section hold beyond 0.4, above bounds of 0.32 and 0.29; the tumor axis was not tested because the core contains no tumor. Two of the three findings hold beyond the bound of 0.38 on Lung FFPE, but the tumor axis breaks at 0.4, the first tested value above the bound. The signaling-gene lean and the tumor axis both break at 0.2 on Ovarian FF, below the bound of 0.49. Spillover below the bound is not ruled out for these three findings, which should therefore be treated as possible artifacts. The bound of Ovarian FF also lies above the tested range, so the relocation finding on this section, which holds up to 0.4, is undecided.

**Fig. 5:**
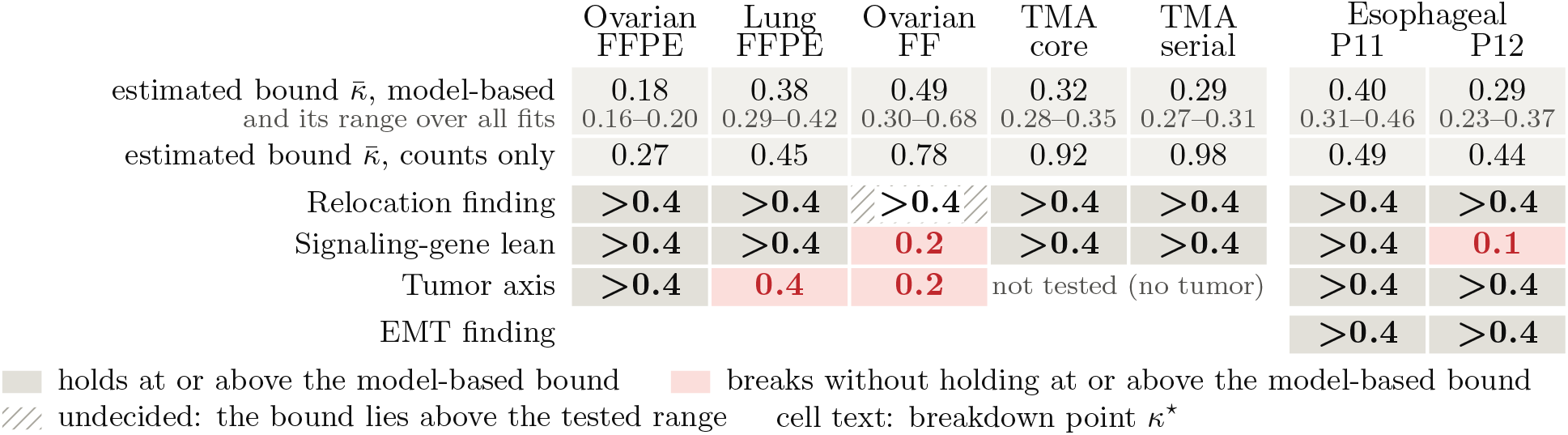
Breakdown point of each finding against the estimated bound on spillover of each section. Each entry is the breakdown point *κ*^*⋆*^ of a finding on a section (*>*0.4 if the finding holds at every tested *κ*). The top rows show the estimated bound 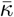, model-based with its range over all fits, and from the counts alone. Colors indicate the outcome against the model-based bound. Gray: the finding holds at a tested *κ* at or above the bound, so spillover within the bound does not explain it. Red: the finding breaks without holding at any tested *κ* at or above the bound, so spillover is not ruled out. Hatched: the finding holds up to 0.4, the largest tested *κ*, but the bound lies above 0.4, so the finding is undecided. The last two columns are the esophageal cancer sections on the published counts, with each bound the highest over three versions of the counts (all versions in Fig. S2).

#### A published finding in esophageal cancer with-stands spillover

Tumor cells of esophageal adenocarcinoma in niches infiltrated by immune or stromal cells express more of an epithelial–mesenchymal transition (EMT) program [45]. Spillover could produce the same difference, since cancer-associated fibroblasts score high on common EMT signatures [49] and transcripts of these fibroblasts can be assigned to tumor cells in stroma-infiltrated niches. We tested this finding on two sections from the study, with the published definitions of the program and of the niches [50]. The EMT finding holds in the clean profiles of DisCell beyond *κ* = 0.4 in both patients, against bounds of 0.36 to 0.40 and 0.27 to 0.29, on the published counts as well as on two recounts of the raw transcripts, so the finding withstands spillover of the modeled kind. The relocation finding and the tumor axis also hold beyond 0.4 in both patients, whereas the signaling-gene lean breaks at 0.1 on the published counts of the second patient (P12), where the response accounts for little of the shift between niches (S5.3).

### 3.3 Relocation predicts a cell type in another niche, with its spillover separated

#### In simulation, the response part recovers the spillover-free shift

We use the niche-dependent prior of the response to predict how the expression of a cell type would change in another niche, which we call relocation, with niches defined as ten clusters of neighbor composition (S3.2). To relocate a cell from one niche to another, the intrinsic state of the cell is kept, the response is set to the response expected for the cell type in the target niche, the spillover influx is replaced by that of the target niche, and the result is decoded. The predicted shift in the mean expression of the type then has a response part and a spillover part (Fig. 6a). The response part explains about three quarters of the clean shift, the true shift without spillover, in tissue simulated from the model, at every true spillover fraction (Fig. 6b). A regression of the observed shifts on neighbor composition, which cannot separate spillover, explains 40% with no spillover and 14% with the highest spillover fraction, on the log scale (S5.1).

**Fig. 6:**
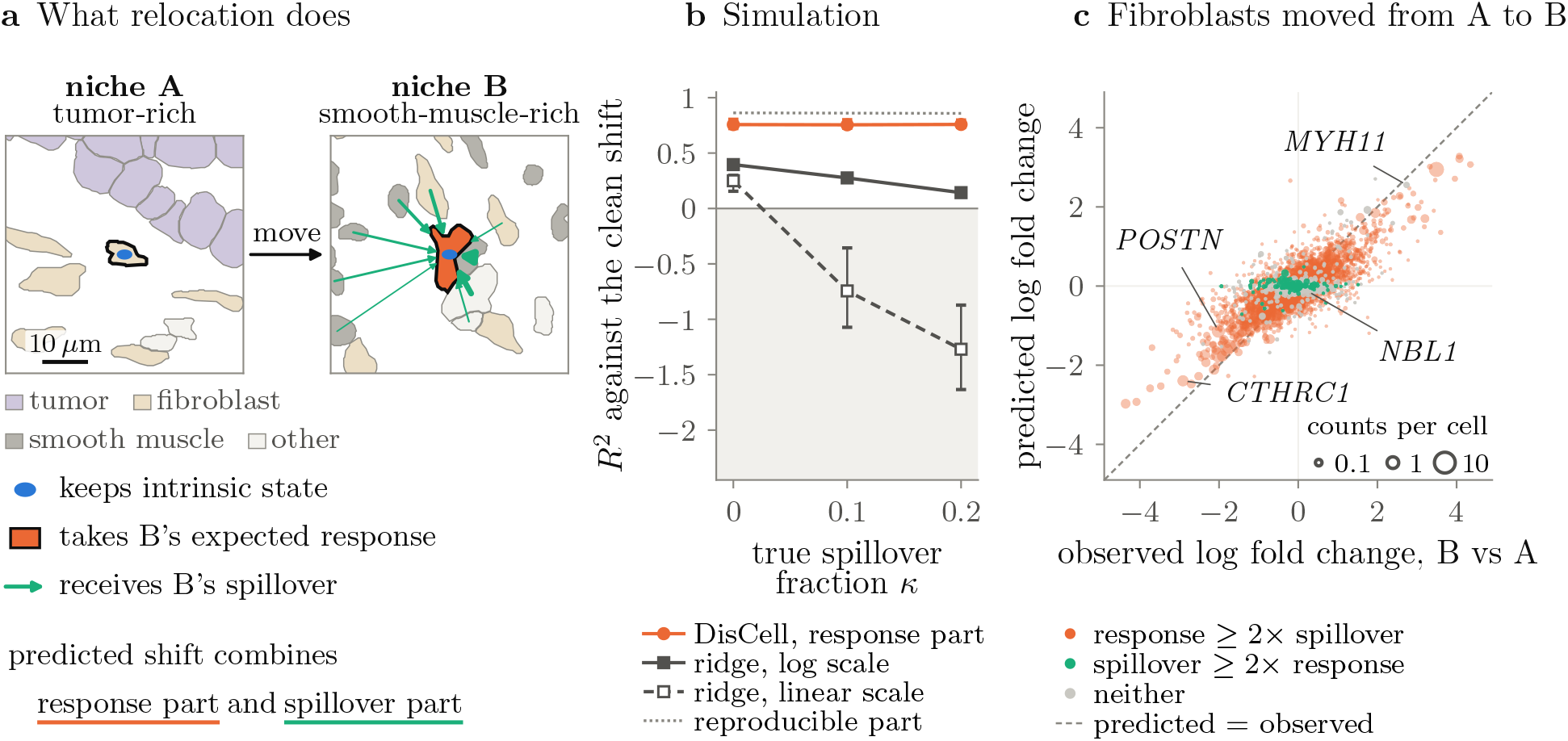
Relocation. **a**, What changes for a moved cell (Sec. 3.3), on segmented cells of Ovarian FFPE around a typical fibroblast of each niche. Arrow widths are each neighbor’s weight in the spillover. **b**, The share of the clean shift, the true shift without spillover, explained in simulation, with the range over three seeds as bars. The ridge regression predicts the shifts from neighbor composition, on the log or the linear scale, and cannot separate spillover. The shaded region is worse than predicting no change, and the dotted line is the reproducible part of the clean shift. **c**, Fibroblasts moved between the two niches of a at *κ* = 0.1, in one fit. Each point is a gene, placed by the observed and the predicted log fold change of the fibroblasts’ mean expression between the niches, centered over genes. Genes are colored when one part is at least twice the other, and gray otherwise, and dot area follows the gene’s mean counts per cell in fibroblasts.

#### On withheld cells, DisCell predicts with lower error than Cellina on most tasks and says how much of the change is spillover

Most of the predicted shift for fibroblasts moved from a tumor-rich to a smooth-muscle-rich niche of Ovarian FFPE is in the response part, as for *CTHRC1* (Fig. 4), while *NBL1*, an intrinsic gene in Fig. 3, barely changes, and for the smooth-muscle gene *MYH11* the spillover part is about as large (Fig. 6c). We then compared DisCell with Cellina, the competing method designed for this prediction, following the protocol of its benchmark [28] (Fig. 7, S3.3). We defined nine tasks on the four fitted sections, in each of which all cells of one type in one niche were removed before fitting and predicted from cells of the same type in another niche. DisCell and Cellina were each fitted with three seeds, DisCell with the settings used throughout and at *κ* = 0.1, set before the test. The error of DisCell is lower than that of Cellina as published on eight of the nine tasks, the exception, the tumor cells of Ovarian FF, being within the spread over seeds, and the fold changes predicted by DisCell correlate better with those of the withheld cells on all nine. The error of Cellina decreases on seven to nine tasks when it is given the same inputs as DisCell, and DisCell still has the lower error on six. The best form of Cellina for each task has a lower error than DisCell on four tasks, but this form differs between tasks and choosing it requires the withheld cells. A ridge regression of the shifts on neighbor composition has the lowest error on five tasks, but it involves no relocation of cells, its predicted shift includes spillover, and it explains much less of the simulated clean shift than the response part of DisCell (Fig. 6b).

**Fig. 7:**
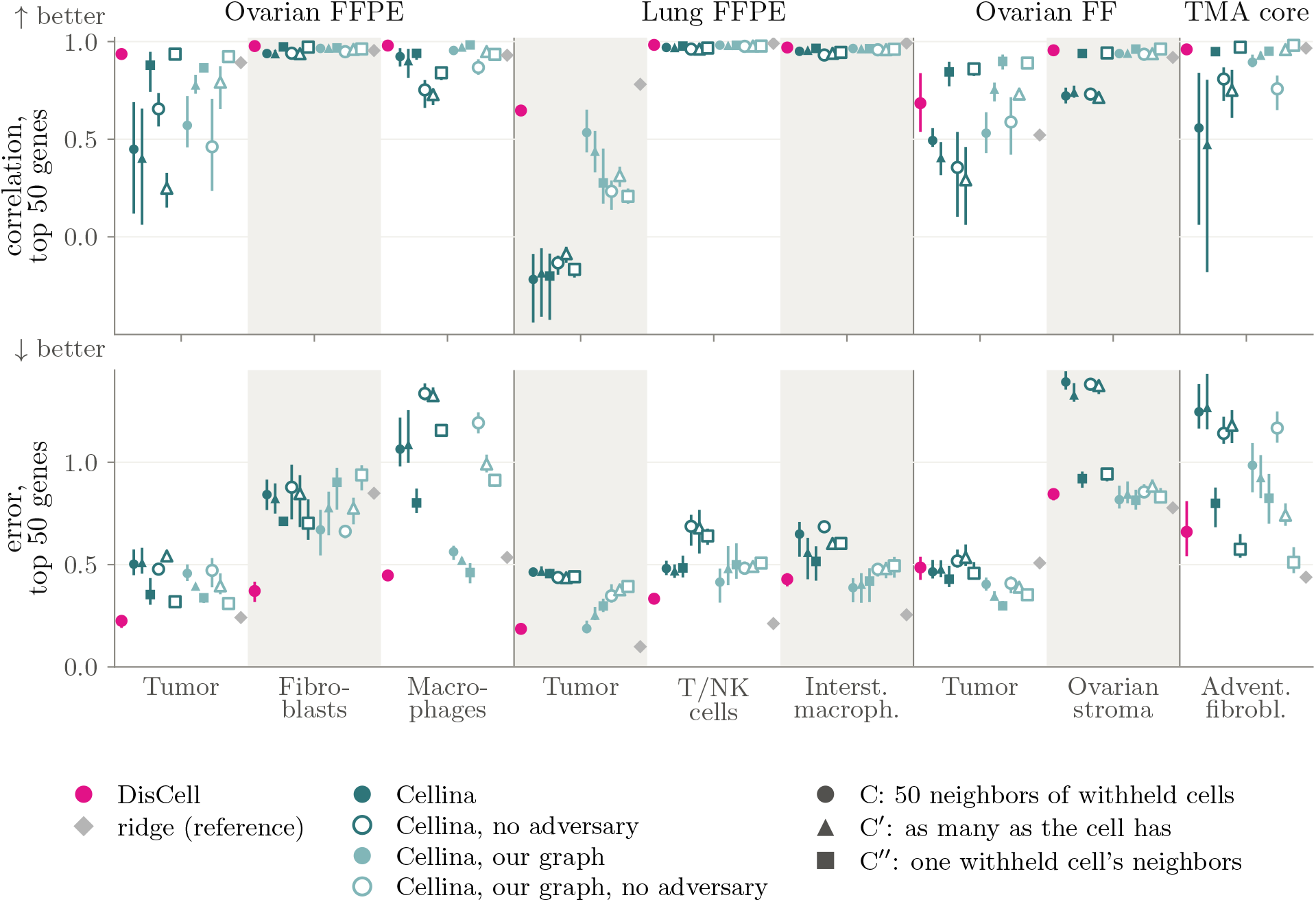
Relocation of withheld cells. In each task, all cells of one type in one niche were removed before fitting and predicted from cells of the same type in another niche (S3.3). Correlation (top) and error (bottom) of the predicted log fold changes on the 50 most changed genes; means over three seeds, with lines spanning the range. Cellina as published (filled dark teal circles) uses its own cell graph, its domain adversary and 50 pooled neighbors of the withheld cells; the other forms use our cell graph (light teal), as many neighbors as the cell has (triangles) or the neighbors of one withheld cell (squares), each also without the adversary (hollow). The ridge regression (gray) is a reference that cannot separate spillover. Values are in Tables S5–S7.

## 4 Discussion

We have introduced DisCell, a generative model of spatially resolved single-cell transcriptomics data that evaluates the impact of spillover in downstream analyses. The technical effect of transcript spillover and the biological effect of niche response both make the observed expression profiles depend on their surroundings. To address the effect of this confounding in biological interpretations, DisCell tests whether a finding computed from the data, such as a niche program, is robust to spillover effects. The workflow fits the model over a range of spillover fractions and treats a finding that breaks before the section’s estimated bound as possibly an artifact of spillover. One fit takes under ten minutes for 400,000 cells (Table S3), so the test can be added to an existing analysis at little cost.

On the sections studied here, no competing method kept more of the cell cycle in its intrinsic state while holding less of the neighborhood. Most of the findings we tested were robust to spillover, including a published association between EMT and infiltrated niches in esophageal cancer, while three findings were flagged as potential artifacts, which a single fit at one spillover fraction would not have shown. In relocation experiments, DisCell predicted single-cell expression profiles withheld from training with lower error than Cellina in most tasks, and separated the spillover contribution of each change.

### Limitations and outlook

The test covers only the spillover modeled as one fraction per section, shared across genes, and reaching only one ring of neighbors. It also cannot separate a response from intrinsic variation that is organized in space, such as a cancer subclone occupying one region [51], which is read as response. Additionally, extending the model to allow cell-specific spillover effects could turn the upper bound into an estimate. Because DisCell is fitted to one section at a time, it needs no batch correction. Modeling batch effects would allow a single fit across many sections, donors and platforms. Section S1 discusses these and further limitations. Overall, we expect DisCell and future extensions to increase the reproducibility of reported biological discoveries from large-scale spatial single-cell data analysis studies.

## Acknowledgments

This work was partially supported by the Wallenberg AI, Autonomous Systems and Software Program (WASP) funded by the Knut and Alice Wallenberg Foundation.

