## Supplemental for "DisCell: Separating Spillover from Niche Response in Spatial Transcriptomics"

### DisCell: Supplementary Material

#### S1 Limitations and future work

**Limitations.** The bound and the breakdown points cover spillover of the modeled kind only. It is one fraction per section, the same for every gene, and it reaches one ring of neighbors. The bound differs between the cell types of a section, and we report the loosest (Sec. S2.2). Spillover that is specific to some genes [1], as ambient contamination is in droplet data [2], or that spares the nucleus, is not covered, and whatever part of it the model cannot explain may be absorbed by the response. The test also cannot separate a response from intrinsic variation that is organized in space, such as a subclone filling one region [3] or a state that follows a gradient across the tissue [4, 5], or from cells that select their niche. Because the intrinsic state is kept free of the niche, such variation ends up in the response. Responses to direct contact act between adjacent cells [6], the same short range as spillover, which makes them the hardest to separate from it. The cell-type labels are taken as given, although spillover can also corrupt them, and a finding that rests on cell states or niche labels derived from expression would need them recomputed on the clean profiles, which we do not do here. At the weights used here, the response describes how a cell type shifts with its niche rather than how one cell deviates from it, and a per-cell response built into a simulation was not detected. Relocation therefore predicts a type's expression in another niche, not the effect of moving one cell. The simulations in which the breakdown point separates spillover from response follow the model's own form of spillover (Sec. S5.1). All sections come from one platform. The model needs only counts, positions and type labels, and the image is optional, so it applies to other platforms with segmented cells, imaging-based ones such as MERSCOPE or CosMx and sequencing-based ones such as Visium HD, but this is untested.

**What could be improved.** A second view of each cell at a different spillover level, such as the split of its transcripts between nucleus and cytoplasm, could turn the upper bound on $\kappa$ into an estimate, as nuclear counts already identify other artifacts [7], although the nucleus also receives some spillover [1]. Gene-specific spillover fractions would cover the cases the single fraction misses. A weaker pull of the response toward its prior would open the per-cell channel, at some cost to how well the intrinsic state keeps cell identity. The separation itself could improve with a richer description of the niche, for example a learned one [8] or one over several scales [9]. The response's prior depends on the niche descriptor, and without an informative prior the condition-specific representation collapses into the shared one [10], so a descriptor that captures more of the niche gives the response more of it to hold. The adversary removes only the niche information it is asked to predict, here the composition and an image niche, so richer targets would remove more of it from the intrinsic state.

#### S2 Theory

##### S2.1 The objective

Fix a cell $i$ and hold its neighbors' clean profiles, and hence $\bar{\boldsymbol{\rho}}_i$, fixed. The counts are multinomial, so $\ln p(\mathbf{x}_i \mid \mathbf{z}_i, \mathbf{w}_i) = \ell_i \, \mathrm{Rec}_i(\mathbf{w}_i) + \text{const}$, where the constant is the logarithm of the multinomial coefficient and depends on the counts alone. With the two posteriors of Sec. 2.2, Jensen's inequality gives the evidence lower bound

$$\ln p(\mathbf{x}_i \mid \mathbf{c}_i, t_i) \geq \mathbb{E}_{q(\mathbf{z}_i \mid \mathbf{x}_i, t_i)} \Big[ \ell_i \, \mathbb{E}_{q(\mathbf{w}_i \mid \mathbf{c}_i, t_i, \mathbf{z}_i, \mathbf{x}_i)} \, \mathrm{Rec}_i(\mathbf{w}_i) - \mathrm{KL}\left( q(\mathbf{w}_i \mid \mathbf{c}_i, t_i, \mathbf{z}_i, \mathbf{x}_i) \,\|\, p(\mathbf{w}_i \mid \mathbf{c}_i, t_i) \right) \Big]$$
$$- \mathrm{KL}\left( q(\mathbf{z}_i \mid \mathbf{x}_i, t_i) \,\|\, p(\mathbf{z}_i) \right) + \text{const} =: L_i.$$

The divergence of the response sits inside the expectation over $\mathbf{z}_i$ because its posterior depends on $\mathbf{z}_i$, and training evaluates both expectations at one reparameterized draw. Using the prior $p(\mathbf{w}_i \mid \mathbf{c}_i, t_i)$ in place of the response's posterior gives a second bound on the same evidence, without the divergence of the response,

$$\ln p(\mathbf{x}_i \mid \mathbf{c}_i, t_i) \geq \ell_i \, \mathbb{E}_{q(\mathbf{z}_i|\mathbf{x}_i,t_i)} \, \mathbb{E}_{p(\breve{\mathbf{w}}_i|\mathbf{c}_i,t_i)} \, \mathrm{Rec}_i(\breve{\mathbf{w}}_i) - \mathrm{KL}\left(q(\mathbf{z}_i \mid \mathbf{x}_i, t_i) \,\|\, p(\mathbf{z}_i)\right) + \mathrm{const} =: L_i^z,$$

and its reconstruction term is the second reconstruction of Sec. 2.2. Each bound carries its own divergence of the intrinsic state, so $L_i + \omega L_i^z$ contains it $1 + \omega$ times. Let $\bar{\ell}$ be the section's median total count, except on Ovarian FFPE, where it is 143 against a median of 178. For a cell with $\ell_i = \bar{\ell}$, $(L_i + \omega L_i^z)/\bar{\ell}$ is, up to a constant, the expectation of the bracket of Eq. (7) at $\alpha_a = 0$ and $\alpha_z = \alpha_w = 1/\bar{\ell}$, with $\mathbf{z}_i$ and $\mathbf{w}_i$ drawn from their posteriors and $\breve{\mathbf{w}}_i$ from the prior. Dividing the reconstruction by each cell's own total instead of by $\bar{\ell}$ lets the weights carry over between sections. The second path of Slavutsky et al. [10] decodes the invariant latent alone with its own decoder, while here both paths share one decoder and bound the same evidence. We use $\tau = 20\,\mu$m, $\omega = 1$, $\alpha_z = \frac{1}{2}/\bar{\ell}$ (rounded), $\alpha_w = 0.1$, $\alpha_a = 0.3$ and $\lambda_y = 3$. The weight $\alpha_w$ rises linearly from zero over the first 30 epochs, and no checkpoint is kept before then. At these weights $J$ is a weighted surrogate of the bound. The bounds of different cells are coupled, since $\bar{\boldsymbol{\rho}}_i$ depends on the neighbors' latents. Neighbors enter each training step through one draw from their own posteriors and without gradient (Sec. S4), so training treats the current $\bar{\boldsymbol{\rho}}_i$ as data. With the adversary, training seeks a saddle point rather than a maximum of $J$, so we claim no likelihood-based uncertainty, and the intervals of Sec. 2.4 come from the seeds and from resampling tissue.

#### S2.2 Identification

*Proof of Proposition 1.* With a neighbor, $\sum_g \bar{\rho}_{ig} = 1$, so the clean profile implied by $\kappa'$, $\boldsymbol{\rho}_i(\kappa') = (\mathbf{p}_i - \kappa'\bar{\boldsymbol{\rho}}_i)/(1-\kappa')$, sums to one. Its share of gene $g$ is non-negative if and only if $\kappa'\bar{\rho}_{ig} \leq p_{ig}$, which holds for every gene if and only if $\kappa' \leq \bar{\kappa}_i$. Since $p_{ig}/\bar{\rho}_{ig} = \kappa + (1-\kappa)\rho_{ig}/\bar{\rho}_{ig} \geq \kappa$, the true $\kappa$ satisfies the bound. □

One cell's bound is useless, with a median over a type's cells of 0.6 to 3, so we pool over the connected cells $C_t$ of type $t$,

$$X_{tg} = \textstyle\sum_{i \in C_t} x_{ig}, \qquad S_{tg} = \sum_{i \in C_t} \ell_i \, \bar{\rho}_{ig}, \qquad \bar{\kappa}_t = \min_{g:\, S_{tg} > 0} \mathbb{E}\, X_{tg}/S_{tg}. \tag{S1}$$

**Proposition S1.** *Under the model of Sec. 2.1, with the neighbors' clean profiles fixed, $\kappa \leq \bar{\kappa}_t$ for every type $t$, and $\bar{\kappa}_t \geq \min_{i \in C_t} \bar{\kappa}_i$.*

*Proof.* By Proposition 1, $\kappa \ell_i \bar{\rho}_{ig} \leq \ell_i p_{ig}$ for every connected cell and gene, and summing over $C_t$ gives $\kappa S_{tg} \leq \mathbb{E}\, X_{tg}$. A ratio of sums of positive terms is at least its smallest term ratio, hence the second part. □

The expected count in Eq. (S1) is replaced by the exact Poisson upper limit $U_a(n) = \frac{1}{2}\chi^2_{1-a,\,2(n+1)}$ [11], valid for any count, at $a = 0.05/G_t$. This Bonferroni correction over the $G_t$ genes with $S_{tg} \geq 25$, which would receive at least 10 spilled counts at $\kappa = 0.4$, keeps the minimum from being set by chance. The *model-based* estimate computes each type's bound from each of the three fits at $\kappa = 0.1$ and averages the three, and the *counts-only* estimate takes the neighbors' observed profiles, with a lower confidence limit on $S_{tg}$. Since a section-wide $\kappa$ must respect every type (Proposition S1), the tightest valid section bound is the smallest type estimate, 0.04 to 0.13 per section. We report the largest, $\bar{\kappa} = \max_t \bar{\kappa}_t$, which is the loosest valid choice, so a finding that holds beyond it holds beyond every type's estimate. Sec. S5.2 gives its range over all fits of a section. The minimum is set by a few genes where the model's form of spillover fits worst, markers of small, low-count neighbor types such as *CD8A*, whose spillover, scaled by the receiving cell's total count, exceeds what the neighbors hold, and nuclear-retained RNAs such as *MEG3*, and on that misfit it would place the value 0.1 used for single-$\kappa$ results above the bound on four of the five sections. Both versions bound spillover of the model's form only.

### S3 Definitions

#### S3.1 Neighborhood information and the cell-cycle $R^2$

**Neighborhood information.** The probe predicts the neighbor composition $\mathbf{y}_i$ from $(\boldsymbol{\mu}_{z,i}, t_i)$. Fitted on training tiles and scored on held-out tiles, its gain

$$\Gamma = \frac{1}{K} \sum_{k=1}^{K} \frac{1}{2} \ln \frac{\mathbb{E}\left(y_{ik} - \bar{y}_k(t)\right)^2}{\mathbb{E}\left(y_{ik} - \hat{y}_{ik}\right)^2},$$

with $K$ the number of cell types, $y_{ik}$ the share of type $k$ among cell $i$'s neighbors, $\bar{y}_k(t)$ its mean over cells of type $t$ and $\hat{y}_{ik}$ the prediction, estimates the predictive information from $\mathbf{z}$ to the composition given the type [12]. The neighborhood information $1 - e^{-2(\Gamma - \Gamma^{\text{perm}})}$, with $\Gamma^{\text{perm}}$ the gain after permuting $\boldsymbol{\mu}_z$ within types, approximates the share of within-type variation recovered beyond chance, here by a perceptron. The ridge probe is the same with a ridge regression in place of the perceptron.

**Cell-cycle $R^2$.** A separate ridge regression, not the probe, measures how much of the cell cycle a latent keeps. It regresses the S and G2/M scores [13, 14], centered per type, on the latent. The cells are the tenth of held-out cells with the highest sum of the two scores, unassigned cells excluded, together with the training cells that score at least as high. In other cells the scores are mostly noise, and the selection uses the scores alone, not the type labels. The ridge is fitted on the training cells and scored on the held-out ones, which gives the cell-cycle $R^2$. Spillover enters the scores, so the $R^2$ shows where the model places the cycle.

**NMI.** The normalized mutual information compares a $k$-means clustering of $\boldsymbol{\mu}_z$ with the type labels.

#### S3.2 Niche pairs and their shifts

Niches are ten $k$-means clusters of neighbor composition, and tiles are dealt into five spatial folds, one scored and four supplying the model quantities. The folds split the readout, not the training, so most scored cells lie in tiles the models were fitted on. A niche pair is a type $t$ and niches $A$ and $B$, each with at least 500 model-side and 100 scored cells of $t$. With $\boldsymbol{\lambda}_N$ the mean observed shares of the scored cells in niche $N$, and $\mathbf{m}_N$, $\boldsymbol{\rho}_N$ and $\bar{\boldsymbol{\rho}}_N$ the model-side means of $m(\mathbf{c}_i, t)$, $\boldsymbol{\rho}_i$ and $\bar{\boldsymbol{\rho}}_i$, per gene

$$\Delta_{\text{obs}} = \ln \boldsymbol{\lambda}_B - \ln \boldsymbol{\lambda}_A, \quad \Delta_{\text{resp}} = \mathbf{B}(\mathbf{m}_B - \mathbf{m}_A), \quad \Delta_{\text{spill}} = \ln \tfrac{(1-\kappa)\boldsymbol{\rho}_A + \kappa\bar{\boldsymbol{\rho}}_B}{(1-\kappa)\boldsymbol{\rho}_A + \kappa\bar{\boldsymbol{\rho}}_A}. \tag{S2}$$

DisCell's predicted shift is $\ln \frac{(1-\kappa)\boldsymbol{\rho}_{A\to B} + \kappa\bar{\boldsymbol{\rho}}_B}{(1-\kappa)\boldsymbol{\rho}_A + \kappa\bar{\boldsymbol{\rho}}_A}$, with $\boldsymbol{\rho}_{A\to B} \propto \boldsymbol{\rho}_A \exp(\Delta_{\text{resp}})$. A pair's noise ceiling is the Spearman–Brown reliability of $\Delta_{\text{obs}}$, centered over genes, from two random halves of the scored cells. It measures how reproducible the difference between the two niches is, and Fig. 6c shows the fibroblast pair with the highest noise ceiling. The ridge regression fits the expression of the model-side cells of $t$, on the linear or the log scale, on their neighbor composition, with the penalty set by generalized cross-validation, and predicts the shift from the compositions of the two niches.

#### S3.3 Relocation of withheld cells

This benchmark, Fig. 7, follows the leave-one-out protocol of Cellina [15], the competing method built for this prediction. resolVI has no niche term and SIMVI estimates an average niche effect rather than moving cells (Table 1). MintFlow's software has no protocol for predicting a cell type moved to another niche, and the adaptation in Cellina's benchmark fits it on the withheld cells, which this test forbids. A task is a cell type $t$, a source niche $A$ and a target niche $B$, taken from the niche pairs of Sec. S3.2 with at least 500 cells of $t$ in each niche and a noise ceiling of at least 0.5. Pairs are ranked by their number of target cells, and each section contributes its first pair for each new type, up to three tasks per section and two on Ovarian FF, which gives nine tasks. All cells of $t$ in $B$ are removed from the data before fitting, with their edges, so that no model sees their expression. Their positions are used at prediction, as in Cellina's protocol. DisCell is refitted at the final settings at $\kappa = 0$, 0.1 and 0.4, with three seeds except at $\kappa = 0$ and 0.4 on Ovarian

FF, which have one, and Cellina at its published settings in the twelve forms below, each with three seeds. To move a cell of $t$ from $A$, DisCell computes the niche descriptor and the spillover influx at each withheld cell's position from its remaining neighbors and its image, averages them over the withheld cells, and decodes the cell at its own $\boldsymbol{\mu}_z$ with the resulting response and influx. The image at a withheld cell has the cell itself masked out but shows the cells around it, which include other withheld cells, so DisCell sees their appearance, though never their expression. Cellina uses no image. As published, Cellina is fitted on its own cell graph, which links each cell to up to 200 neighbors with Gaussian weights of bandwidth 100 $\mu$m, with its domain adversary on the annotated regions, here the niche labels that define $A$ and $B$. Its form C replaces a moved cell's neighbors with 50 cells drawn from the neighbors of the withheld cells that are not of type $t$. Their log-normalized expression is averaged, as in Cellina's training features. Cellina's released benchmark script averages raw counts here, which its later code corrects. The other forms give Cellina the inputs DisCell has. On our graph (ours) it sees the same neighbors as DisCell. C′ draws as many cells as the moved cell has neighbors, about six on our graph, because fifty lies far outside the neighborhoods Cellina saw in training there. C″ gives each moved cell the neighborhood of one withheld cell drawn at random, the information DisCell has at prediction. It is the anchored counterfactual that Cellina's current code makes its default, run with that code's own function and the settings its documentation gives for leave-one-out. The neighborhood is the mean over the withheld cell's neighbors on the full section, weighted as in training, with every cell of type $t$ left out, and withheld cells left without a neighbor are skipped. This code is commit dd860d0 of Cellina's repository, version 1.2.0 of its changelog, dated 1 October 2026 and not yet a tagged release. It changes only the counterfactual functions of the release 1.1.1 every Cellina model here was fitted with, and loaded under it each model reproduces its earlier predictions exactly. A star marks a form fitted without the adversary, and every form ran on both graphs (Tables S5 and S6). The ridge regression of Sec. S3.2, fitted to the shares of the remaining cells of $t$, shifts each cell's observed expression proportions by its predicted shift from the mean composition of $A$ to the mean composition around the withheld cells, and renormalizes (Table S7). The withheld cells are the truth and the observed cells of $t$ in $A$ the control. Expression is scaled to $10^4$ counts per cell and averaged over cells, and the log fold change of a gene is $\ln(\bar{p}_{\text{pert}} + 1) - \ln(\bar{p}_{\text{ctrl}} + 1)$, with the moved cells as the perturbed mean for the prediction and the withheld cells for the observation. The 50 genes are those with the largest observed fold change between the control and the withheld cells, the rule of Cellina's benchmark, and the scores are the Pearson correlation and the root mean squared error between predicted and observed fold changes over these genes.

#### S3.4 The findings and their intervals

A finding is a single number $r$, computed from one fitted model, that is zero in expectation when the effect is absent, positive in the direction the finding claims and negative in the opposite one. The breakdown point uses only the sign of $r$, so $r$ need not grow in proportion to the effect, but it must not be positive by construction, as a squared or absolute effect on its own would be. Table S1 lists the findings tested and how each $r$ is computed. To test a finding at a spillover fraction $\kappa$, the model is fitted at $\kappa$ with three seeds and $r$ is computed on each fit. Only what the fitted model supplies changes with $\kappa$: the clean profiles $\boldsymbol{\rho}_i$, the latents and the split of a predicted shift into response and spillover. The cells, labels, niches, niche pairs and gene sets that enter $r$ stay fixed. The result at $\kappa = 0$ is what an analysis that ignores spillover would report, not a confirmation of the claim. A finding that does not hold there is no finding and has no $\kappa^\star$.

The interval $\mathrm{CI}_\kappa(r)$ of Definition 1 measures how much $r$ varies over the tissue and between fits. Within each fit, the 200 $\mu$m squares are resampled with replacement or, for the findings marked † in Table S1, half of them are drawn without replacement, and $r$ is recomputed on each draw. The cycle and niche-information checks and the esophageal score form use 10,000 draws, and the other findings 4,000. Each draw of the interval then picks three of the fits with replacement, takes an independent draw of squares for each fit picked and averages them, so that it covers the spread between fits as well as over the tissue. $\mathrm{CI}_\kappa(r)$ runs from the $\alpha/(2M)$ to the $1 - \alpha/(2M)$ quantile of these averages, with $\alpha = 0.05$ and $M$ the number of findings tested on the section, so that all findings of a section are covered together. A finding holds at $\kappa$ if $\mathrm{CI}_\kappa(r)$ lies entirely above zero. With three fits, one fit near zero can widen the interval enough to reach zero, which is how both tumor-axis breaks arise, and more fits narrow it rather than adding chances to fail.

The correction counts $M = 7$ findings on Ovarian FFPE, Lung FFPE and Ovarian FF, 6 on the TMA sections and 13 on the esophageal ones, which add the six program findings. These counts include two tested

Table S1: The findings tested by the breakdown point. The first three rows and the EMT finding appear in Fig. 5. Tildes mark centering over a niche pair's genes, $A$ and $B$ are the two niches of a pair or, on the esophageal sections, the published niche groups, the shifts $\Delta$ and means $\mathbf{m}_N$ are those of Eq. (S2), and † marks the findings whose interval resamples half of the squares without replacement.

| Finding | $r$ | $r > 0$ means |
|---|---|---|
| Relocation finding† | mean over niche pairs of $2\langle\tilde{\Delta}_{\text{obs}} - \tilde{\Delta}_{\text{spill}}, \tilde{\Delta}_{\text{resp}}\rangle / \|\tilde{\Delta}_{\text{obs}}\|^2$ | the response points along the part of the observed shift that spillover leaves |
| Signaling-gene lean† | rank-biserial effect, for the ligand and receptor genes of CellChat [16] against the other genes, of each gene's response share $\sum \lvert\tilde{\Delta}_{\text{resp}}\rvert / \sum \big(\lvert\tilde{\Delta}_{\text{resp}}\rvert + \lvert\tilde{\Delta}_{\text{spill}}\rvert + \lvert\tilde{\Delta}_{\text{obs}} - \tilde{\Delta}_{\text{resp}} - \tilde{\Delta}_{\text{spill}}\rvert\big)$, with sums over niche pairs | ligand and receptor genes take a larger share of the response than other genes |
| Tumor axis | mean over type–gene pairs of the absolute Kendall correlation of $\mathbf{Bm}_N$ over bands $N$ of the local tumor share, minus that over bins of the in-plane coordinate least correlated with the bands (correlation 0.41 on Ovarian FFPE) | the response follows the tumor more than a plain tissue coordinate; not on the TMA sections, which lack tumor cells |
| Cycle check | cell-cycle $R^2$ of $\boldsymbol{\mu}_z$ minus that of $\boldsymbol{\mu}_w$ (Sec. S3.1) | the cell cycle sits in the intrinsic state rather than the response |
| Niche-information check | $\text{MI}(\boldsymbol{\mu}_w; N) - \text{MI}(\boldsymbol{\mu}_w; N')$, with $N$ the niche and $N'$ the niche permuted within types [17] | the response carries the niche |
| Esophageal program findings | score form: mean over $B$ minus mean over $A$ of the mean over $g \in \mathcal{G}$ of $\ln(1+10^4\rho_{ig})$, with $\mathcal{G}$ a published program's genes in the panel, each gene standardized over the tumor cells of held-out tiles; response form†: $2\langle\tilde{\Delta}_{\text{obs}} - \tilde{\Delta}_{\text{spill}}, \mathbf{u}\rangle\langle\tilde{\Delta}_{\text{resp}}, \mathbf{u}\rangle / \langle\tilde{\Delta}_{\text{obs}}, \mathbf{u}\rangle^2$, with $\mathbf{u}$ the indicator of $\mathcal{G}$ | the program is higher in tumor cells of $B$ than of $A$; the EMT finding and two further programs [18], each in both forms |

findings that are not reported, comparisons of moved cells with the type's mean profile and with a random cell. The cycle and niche-information checks hold beyond 0.4 on every section.

#### S3.5 The per-gene split

Setting each of the decoder's inputs $\mathbf{z}_i$, $\mathbf{w}_i$ and $\bar{\boldsymbol{\rho}}_i$ to the cell's value or the type's mean, in all eight combinations, gives exact Shapley values $\phi_z$, $\phi_w$ and $\phi_{\bar{\rho}}$ of $v = \ln p_g$, with $\phi_{\bar{\rho}} = 0$ at $\kappa = 0$. Contribution $k$'s share of the within-type variation is $s_k = \text{Cov}_t(\phi_k, v)/\,\text{Var}_t(v)$ over the type's held-out cells, the shares sum to one, and intervals come from a bootstrap over 200 $\mu$m squares. The split is of the model's expected expression $\ln p_g$, not of the observed counts, whose variation within a type is mostly counting noise. For the median gene of a type, $\ln p_g$ explains 1 to 8% of the within-type variation of the observed $\ln(1 + 10^4 x_g/\ell_i)$ on the first seed. For counts drawn from the model itself it explains about as much, so even a correct model would explain little more at these total counts. The genes of each type with at least 300 held-out cells, fixed in advance, are those detected in at least 10% of its cells and the spillover-test genes, below 10% of the share of the type expressing them most and with a neighbor share at least twice their own. A fit assigns the gene to contribution $k$ when the interval of $s_k$ lies above 0.5 and $s_k$ is at least twice each of the other two, the consensus needs two of three seeds, and $\kappa^\star$ is the smallest of 0.2, 0.3 and 0.4 at which the consensus assignment changes. Every assignment to the response or to spillover in Fig. 3 holds at the first tested value at or above $\bar{\kappa}$, and its four types were fixed in advance.

In simulation at the true $\kappa$, recovered and true shares correlate at 0.94 to 0.99, and a $\kappa$ set 0.1 too high adds about 0.1 to the spillover share. On Lung FFPE and the TMA core, the genes assigned to spillover are again markers of neighboring cell types, as on Ovarian FFPE (Fig. S1).

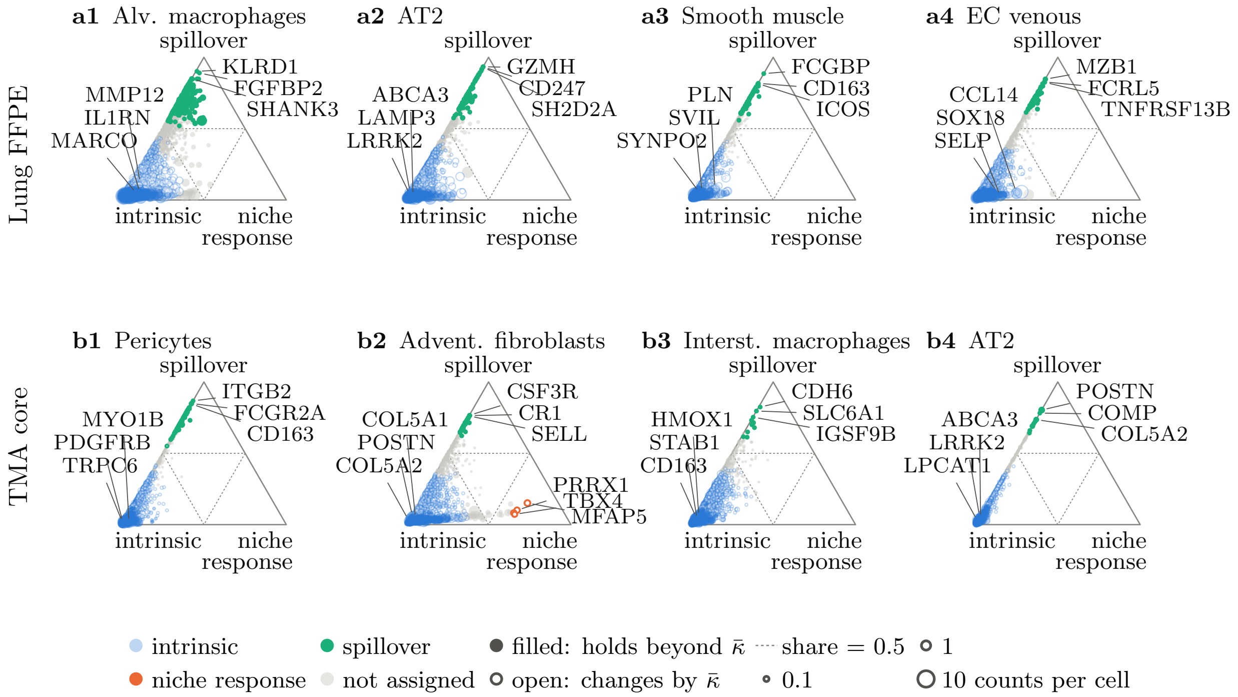


Fig. S1: The per-gene split on two other sections at $\kappa = 0.1$, drawn as in Fig. 3. Each row is one section, with the four cell types with the most genes assigned to spillover. Filled points hold beyond the section's estimated bound $\bar{\kappa}$, 0.38 on Lung FFPE and 0.32 on the TMA core.

### S4 Setup

Table S2: Cells, median counts per cell and cell types, unassigned cells included. The serial section is never fitted, and the esophageal sections have no image.

| | Ovarian FFPE | Lung FFPE | Ovarian FF | TMA core | TMA serial | Esophageal P11 | Esophageal P12 |
|---|---|---|---|---|---|---|---|
| Cells | 407,120 | 278,324 | 1,157,637 | 69,422 | 70,757 | 141,505 | 69,853 |
| Median counts | 178 | 242 | 1,401 | 279 | 253 | 205 | 312 |
| Cell types | 12 | 20 | 10 | 30 | 30 | 7 | 6 |

**Labels, graph and niche descriptor.** Table S2 gives each section's cells, median counts and cell types. Ovarian FFPE's 18 cell types are the curated annotation released with it [19]. On Lung FFPE and Ovarian FF, the 32 and 38 graph-based clusters released with the data were named by scores of marker genes on the panel and mapped to 20 and 10 lineages, and a cluster with no lineage signal was set to unassigned. On the TMA, we clustered both slides jointly into 44 clusters and named each by the correlation of its pooled profile with two lung reference atlases [20, 21], using marker genes for clusters the atlases did not match, such as cartilage and skeletal muscle, which gave 35 curated cell types. Labels are lineages, mapped before any refit

Table S3: Model size and training cost on one GPU, for whole sections. Run time is one whole fit, for DisCell with its evaluations and early stop and for the comparison methods their training time, as the range over three seeds. The second and third seeds of the comparison methods ran beside other fits on the same machine, which accounts for much of their spread. Bold marks the fastest fit per section by its mean over seeds. [a]Not run, out of GPU memory on a 24 GB card (SIMVI) or out of host memory (MintFlow on Ovarian FF).

| | Ovarian FFPE | Lung FFPE | Ovarian FF | TMA core |
|---|---|---|---|---|
| *DisCell, model and training details* | | | | |
| parameters (M) | 4.19 | 4.12 | 4.11 | 4.13 |
| peak memory (GiB) | 6.3 | 4.3 | 15.1 | 1.4 |
| epochs to the stop | 105–115 | 80–100 | 175–230 | 85–90 |
| s per epoch, training only ↓ | 1.98 | 1.35 | 5.78 | 0.45 |
| *Run time of one fit (min)* ↓ | | | | |
| DisCell | 8.7–9.3 | **3.2–7.4** | 31.2–43.4 | **1.3** |
| resolVI | 32.8–69.5 | 22.0–47.4 | 105.0–216.0 | 5.5–13.7 |
| Cellina | **3.3–4.6** | 4.7–6.4 | 49.1–55.3 | **1.1–1.5** |
| no adversary | 6.5–7.7 | 11.2–13.7 | **22.3–31.4** | 2.5–3.5 |
| Cellina, our graph | 3.5–7.9 | 3.6–5.4 | 17.9–42.1 | 1.3–1.6 |
| no adversary | 9.9–11.6 | 8.5–10.6 | 24.5–29.1 | 3.0–3.4 |
| SIMVI | not run[a] | not run[a] | not run[a] | 96.4–192.6 |
| MintFlow, 50 epochs | 877.6–1245.0 | 896.7–1022.4 | not run[a] | 308.6–501.6 |

with states and locations merged, so Ovarian FFPE's 18 curated cell types become 12 and the TMA's 35 become 30. Unassigned cells form their own type, never a target. Neighbors share a face of Voronoi cells clipped to disks of 30 $\mu$m radius, with edges beyond 40 $\mu$m pruned. Polygon contact would be a poor spillover kernel, as only 2.7 to 6.7% of edges join touching polygons. The niche descriptor joins an attention layer over the neighbors' types [22] to the KRONOS embedding [23] of a 128 $\mu$m field of the morphology image with the cell masked by a disk of 25 $\mu$m radius. These fits use the first KRONOS release. A refit of every section with the released code and the same image encoder reproduces every breakdown point except the tumor axis on Lung FFPE, which then holds beyond 0.4. The released code embeds images with KRONOS2, and a refit of Ovarian FFPE with it gives the same breakdown point for every finding.

**Niche labels for Cellina's adversary.** Cellina's adversary needs a domain for every cell, which its code takes from annotated regions, such as normal mucosa, adenoma and carcinoma [15]. Our sections have none, so we construct one. Each cell's neighbor composition on our cell graph is clustered into ten niches by $k$-means, the rule that defines the niches of Sec. S3.2, and cells without a neighbor form an eleventh class. For Fig. 2 the clusters are fitted on training cells only, and a held-out section takes the label of the nearest center of the section its model was fitted on. In the relocation benchmark the niches that define each task serve as the domains. These labels are used only as Cellina's domains, and DisCell is never fitted on them.

**Training and calibration.** Sections are cut into tiles of at most 4,096 cells, 15% held out, and each batch carries a two-hop halo of neighbors that enter without gradient, as neighbors do in resolVI [24]. The latents have $d_z = 20$ and $d_w = 6$, and a checkpoint is kept when held-out reconstruction improves while the NMI of $\mathbf{z}$ stays within 90% of its maximum. Every weight was set once, mostly on Ovarian FFPE, and $d_z$ was not tuned.

#### S4.1 Comparison methods

Each method ran at the model and training settings of its paper on whole sections. On the serial section, MintFlow and Cellina are evaluated with their fits to the TMA core, as DisCell is. resolVI [24] was fitted on the serial section itself, because its software refused to apply the core's model to cells it had not been set up with, and SIMVI [25] was too, because its software has no way to apply a fit to a new section. Both therefore saw the serial section's cells in training. On the whole-slide sections, SIMVI, with batches of 2,048 in place of its full batch, ran out of GPU memory (24 GB). MintFlow [26] ran out of host memory on Ovarian FF (an estimated 92 of 125 GB). Cellina [15] ran on its own graph, which links each cell to up to 200 neighbors with Gaussian weights of bandwidth 100 $\mu$m, with the full gene panel, where its paper used 2,000 variable genes, cells with at least 5 counts, and its release 1.1.1. Our sections have no annotated regions, so its domain adversary reads the niche labels above, as in the relocation benchmark, and every form also ran without the adversary. In Fig. 2, *Cellina, our graph* runs it on the cell graph DisCell and SIMVI use. On either graph the adversary moved Cellina's neighborhood information by at most 1.9 points while its NMI and cell-cycle $R^2$ fell on all five sections. The comparison methods were fitted on cells of every tile, which favors them, and DisCell's weights were calibrated on these sections. Table S3 gives the size and cost of each method on one RTX 4090 GPU. Eighteen DisCell fits at their longest times (168, 133 and 24 minutes) cost less than one MintFlow fit on Ovarian FFPE, Lung FFPE and the TMA core, and Cellina was faster than DisCell on Ovarian FFPE, and without its adversary also on Ovarian FF.

### S5 Supporting results

#### S5.1 Simulation

The recovery test has 6,000 cells of eight clustered types, 60 genes, a response linear in neighbor composition and $\kappa = 0$ or 0.2, on three simulated sections with three seeds. At the matched $\kappa$ the NMI of $\boldsymbol{\mu}_z$ with the true type is 0.80 to 0.96, at or above clusters of the counts in 17 of 18 fits, and the first canonical correlation of $\boldsymbol{\mu}_w$ with the true response 0.83 to 0.95. Without the adversary $\boldsymbol{\mu}_z$ explains 0.11 to 0.19 of the within-type variation of composition instead of at most 0.04, and the correlation falls to 0.29 to 0.91. At any assumed $\kappa$ from 0 to 0.4 on sections simulated at 0.2, the NMI stays at 0.79 to 0.97 and the correlation at 0.78 to 0.94. The control with two known findings has 30,000 cells and $\kappa_{\text{true}} = 0$, 0.1 or 0.2. On its 20 genes without a response, the difference made by spillover alone is the response part's alignment with what the clean shift leaves of the observed shift. Predicted to break at the first tested value at or above $\kappa_{\text{true}}$, it is no finding at 0 and breaks at 0.05 and 0.1, while the genuine response holds beyond 0.4 in every simulated section. The response part of a relocation explains $R^2 = 0.76$ of the clean shift at every true fraction, a ridge regression on neighbor composition 0.40, 0.28 and 0.14 on the log scale, and on the linear scale 0.25 with no spillover and below zero with spillover. These simulated sections follow DisCell's own model.

#### S5.2 Tissue

**The separation.** The ridge probe ranks DisCell lowest of all the fits of Table S4 on every section, including where the perceptron ranks MintFlow or Cellina on our graph without its adversary lower, and MintFlow and Cellina on either graph reach a higher NMI wherever they ran, which rewards fidelity to the labels, except Cellina with its adversary on Ovarian FFPE, and on our graph also on Ovarian FF. With the core's fits the serial section gives an NMI of 0.61 against 0.62 and a cell-cycle $R^2$ of 0.49 against 0.50. The TMA core carries little neighborhood information to begin with, and there the adversary removes about a third of it.

**The estimated bound.** Over a section's 18 fits the model-based value ranges from 0.16 to 0.20, 0.29 to 0.42, 0.30 to 0.68, 0.28 to 0.35 and 0.27 to 0.31 on Ovarian FFPE, Lung FFPE, Ovarian FF, the TMA core and the serial section, so the tumor-axis break of Lung FFPE at 0.4 lies at the bound. It is below 0.4 in 82 of the 83 types with at least 200 connected cells, the exception being the unassigned cells of Ovarian FF (0.49), without which that section's value would be 0.31. The counts-only estimate is larger in every type.

#### S5.3 The esophageal cohort

Gates set before any fit required at least 50,000 cells, a median of about 100 counts per cell and published labels supported by their marker genes, and admitted two of six patients [18, 27]. P4 has 22,000 cells and P8 a median of 49 counts. P10 and P13 are borderline in total counts, with medians of 106 and 94, and 34 and 66% of their cells lie in labels whose markers are not enriched. On P11 the 12% of cells in such labels were set to unassigned, and P12 has none. DisCell was fitted without images on the published counts and

Table S4: Values behind Fig. 2, on held-out cells (Sec. S3.1). *Neighborhood information*, that of the intrinsic latent on composition, in %. *Cell-cycle* $R^2$, from the intrinsic latent, with DisCell's response in parentheses. *NMI*, its agreement with the type. DisCell's neighborhood information and that without the adversary are ranges over three seeds, and all other entries are means over three seeds. Each Cellina row gives its values with and without its adversary on the niche labels (Sec. S4). –, not run.

| | Ovarian FFPE | Lung FFPE | Ovarian FF | TMA core | TMA serial |
|---|---|---|---|---|---|
| *Neighborhood information (%)* ↓ | | | | | |
| DisCell, no adversary | 15.5–16.2 | 10.7–12.0 | 27.6–28.0 | 5.5–6.9 | 6.6–7.3 |
| DisCell | 4.0–5.5 | 4.2–5.0 | 4.4–5.0 | 3.5–4.8 | 4.0–4.8 |
| resolVI, SIMVI | 17.3, – | 10.9, – | 24.4, – | 5.4, 5.0 | 7.1, 6.9 |
| MintFlow | 5.2 | 1.5 | – | 1.3 | 0.6 |
| Cellina | 14.5, 14.6 | 8.9, 9.1 | 22.8, 23.4 | 5.5, 4.6 | 7.9, 8.3 |
| Cellina, our graph | 14.2, 14.0 | 8.6, 8.0 | 22.0, 22.6 | 5.5, 3.6 | 7.3, 7.4 |
| *Cell-cycle* $R^2$ ↑ | | | | | |
| DisCell (its response) | 0.519 (.006) | 0.186 (.003) | 0.757 (.001) | 0.497 (.000) | 0.495 (−.001) |
| resolVI, SIMVI | 0.566, – | 0.155, – | 0.551, – | 0.347, 0.600 | 0.355, 0.606 |
| MintFlow | 0.030 | 0.007 | – | 0.085 | 0.083 |
| Cellina | 0.319, 0.572 | 0.179, 0.270 | 0.677, 0.688 | 0.295, 0.379 | 0.285, 0.332 |
| Cellina, our graph | 0.326, 0.576 | 0.167, 0.292 | 0.646, 0.686 | 0.218, 0.308 | 0.194, 0.268 |
| *NMI* | | | | | |
| DisCell | 0.708 | 0.733 | 0.688 | 0.618 | 0.607 |
| resolVI, SIMVI | 0.721, – | 0.838, – | 0.677, – | 0.722, 0.499 | 0.729, 0.485 |
| MintFlow | 0.932 | 0.978 | – | 0.913 | 0.911 |
| Cellina | 0.641, 0.817 | 0.794, 0.943 | 0.706, 0.833 | 0.769, 0.824 | 0.618, 0.640 |
| Cellina, our graph | 0.662, 0.828 | 0.808, 0.948 | 0.652, 0.832 | 0.759, 0.831 | 0.614, 0.644 |

two recounts, with and without our quality filter. Cancer-associated fibroblasts score high on common EMT signatures [28], so spillover from stromal neighbors is the natural alternative explanation of the finding. The published finding [18], a higher score of the epithelial–mesenchymal transition program cNMF3 in tumor cells of the infiltrated CellCharter [9] niches CC2 and CC4 than in the carcinoma niche CC1, reproduces on both patients (+0.150 and +0.145) and on the clean profiles holds beyond 0.4 in all three count versions, against model-based bounds of 0.36 to 0.40 and 0.27 to 0.29 and counts-only bounds of 0.47 to 0.49 and 0.41 to 0.44. With the response held at its type mean as the intrinsic part, the response carries 40 to 44% of the shift on P11 and 4 to 5% on P12 at $\kappa = 0.1$. On P12 the response's own alignment with the shift breaks at 0.05 to 0.2, and the signaling-gene lean breaks at 0.05 or 0.1 or does not hold at $\kappa = 0$, while on P11 both hold beyond 0.4 (Fig. S2). Of the authors' two further programs, cNMF1 in the score form holds beyond 0.4 on P12 but on P11 at most at $\kappa = 0$, and cNMF4 does not hold at $\kappa = 0$ in either form on either patient. The adversary leaves 0.19 to 0.57 (ridge) and 0.32 to 0.76 (perceptron) of the neighborhood information of a fit without it, against at most 0.21 and 0.76 elsewhere, on the TMA core, so the adversary works about as well here as on the weakest of the other sections. The niche label carries 0.17 to 0.21 nats in $\mathbf{z}$ against 0.19 to 0.23 in $\mathbf{w}$, with more in $\mathbf{z}$ than in $\mathbf{w}$ on P12. The invariance is weaker on these sections, which have no images, so the small response share on P12 does not show that its tumor cells differ in their own state.

| | Esophageal P11 | | | Esophageal P12 | | |
|---|---|---|---|---|---|---|
| | published counts | filtered recount | unfiltered recount | published counts | filtered recount | unfiltered recount |
| estimated bound $\bar{\kappa}$, model-based<br>and its range over all fits | 0.36<br>0.31–0.41 | 0.38<br>0.31–0.46 | 0.40<br>0.31–0.43 | 0.29<br>0.23–0.33 | 0.27<br>0.24–0.35 | 0.28<br>0.23–0.37 |
| estimated bound $\bar{\kappa}$, counts only | 0.47 | 0.49 | 0.49 | 0.41 | 0.44 | 0.42 |
| EMT finding, score form | **>0.4** | **>0.4** | **>0.4** | **>0.4** | **>0.4** | **>0.4** |
| EMT finding, response form | **>0.4** | **>0.4** | **>0.4** | **0.1** | **0.05** | **0.2** |
| cNMF1, score form | no finding | no finding | **0.05** | **>0.4** | **>0.4** | **>0.4** |
| cNMF1, response form | no finding | no finding | no finding | **0.1** | **0.1** | **0.2** |
| cNMF4, both forms | no finding | no finding | no finding | no finding | no finding | no finding |
| Signaling-gene lean | **>0.4** | **>0.4** | **>0.4** | **0.1** | **0.05** | no finding |
| Tumor axis | **>0.4** | **>0.4** | **>0.4** | **>0.4** | **>0.4** | **>0.4** |

holds at or above the model-based bound    breaks without holding at or above the model-based bound
no finding: does not hold at $\kappa = 0$    cell text: breakdown point $\kappa^\star$

Fig. S2: Breakdown points on the esophageal sections, drawn as Fig. 5, for each patient in the published counts and in two recounts (filtered, unfiltered), each with its own estimated bound in the header rows. The published finding is the epithelial–mesenchymal transition program cNMF3, in the score form and the response form. cNMF1 and cNMF4 are the authors' further programs, and the signaling-gene lean and the tumor axis are findings of Table S1. Every other finding holds beyond 0.4 on all six. Colors are those of Fig. 5, and a gray cell marks a finding that does not hold at $\kappa = 0$.

Table S5: Correlation of log fold changes on the 50 most changed genes (higher is better) in the relocation of withheld cells (Fig. 7), for DisCell (D) and every form of Cellina, fitted and predicted on Cellina's own graph (top), as in its benchmark, and on ours (bottom, Sec. S3.3). The forms are as in Fig. 7, with a star marking Cellina without its domain adversary. DisCell is the same in both blocks. Values are means over three seeds with the standard deviation in parentheses, and bold marks the best mean of each block.

| Task | D | C | C′ | C′′ | C* | C′* | C′′* |
|---|---|---|---|---|---|---|---|
| **On Cellina's own graph** | | | | | | | |
| *Ovarian FFPE* | | | | | | | |
| Tumor | **0.94** (.02) | 0.45 (.30) | 0.40 (.31) | 0.88 (.12) | 0.66 (.09) | 0.25 (.09) | **0.94** (.02) |
| Fibroblasts | **0.98** (.01) | 0.94 (.02) | 0.94 (.02) | 0.97 (.00) | 0.94 (.01) | 0.94 (.01) | 0.97 (.01) |
| Macrophages | **0.98** (.00) | 0.92 (.05) | 0.90 (.08) | 0.94 (.03) | 0.75 (.08) | 0.73 (.05) | 0.84 (.03) |
| *Lung FFPE* | | | | | | | |
| Tumor | **0.65** (.02) | −0.22 (.19) | −0.18 (.20) | −0.20 (.19) | −0.13 (.05) | −0.09 (.04) | −0.17 (.04) |
| T/NK cells | **0.98** (.00) | 0.97 (.01) | 0.97 (.01) | **0.98** (.00) | 0.96 (.01) | 0.96 (.01) | 0.97 (.01) |
| Interst. macrophages | **0.97** (.01) | 0.95 (.01) | 0.96 (.01) | **0.97** (.01) | 0.93 (.01) | 0.94 (.00) | 0.94 (.01) |
| *Ovarian FF* | | | | | | | |
| Tumor | 0.68 (.15) | 0.49 (.05) | 0.41 (.08) | 0.84 (.07) | 0.36 (.23) | 0.29 (.21) | **0.86** (.03) |
| Ovarian stroma | **0.96** (.00) | 0.72 (.04) | 0.74 (.03) | 0.94 (.01) | 0.73 (.01) | 0.71 (.02) | 0.94 (.01) |
| *TMA core* | | | | | | | |
| Advent. fibroblasts | 0.96 (.01) | 0.56 (.43) | 0.47 (.57) | 0.95 (.00) | 0.81 (.10) | 0.75 (.13) | **0.97** (.01) |
| **On our graph** | | | | | | | |
| *Ovarian FFPE* | | | | | | | |
| Tumor | **0.94** (.02) | 0.57 (.14) | 0.78 (.05) | 0.87 (.00) | 0.46 (.24) | 0.79 (.12) | 0.92 (.02) |
| Fibroblasts | **0.98** (.01) | 0.97 (.01) | 0.97 (.01) | 0.97 (.01) | 0.95 (.01) | 0.96 (.01) | 0.96 (.01) |
| Macrophages | **0.98** (.00) | 0.95 (.02) | 0.97 (.01) | **0.98** (.01) | 0.87 (.04) | 0.95 (.02) | 0.93 (.03) |
| *Lung FFPE* | | | | | | | |
| Tumor | **0.65** (.02) | 0.53 (.11) | 0.44 (.11) | 0.28 (.15) | 0.23 (.08) | 0.31 (.05) | 0.21 (.04) |
| T/NK cells | **0.98** (.00) | **0.98** (.00) | **0.98** (.00) | **0.98** (.00) | **0.98** (.01) | **0.98** (.01) | **0.98** (.00) |
| Interst. macrophages | **0.97** (.01) | 0.96 (.01) | 0.96 (.01) | 0.96 (.01) | 0.96 (.00) | 0.96 (.01) | 0.96 (.01) |
| *Ovarian FF* | | | | | | | |
| Tumor | 0.68 (.15) | 0.53 (.10) | 0.76 (.05) | **0.90** (.04) | 0.59 (.15) | 0.73 (.03) | 0.89 (.01) |
| Ovarian stroma | **0.96** (.00) | 0.94 (.01) | 0.94 (.01) | **0.96** (.01) | 0.94 (.01) | 0.94 (.01) | **0.96** (.01) |
| *TMA core* | | | | | | | |
| Advent. fibroblasts | 0.96 (.01) | 0.89 (.03) | 0.93 (.02) | 0.95 (.01) | 0.76 (.10) | 0.96 (.00) | **0.98** (.00) |

Table S6: Error of log fold changes on the 50 most changed genes (lower is better) in the relocation of withheld cells (Fig. 7), for DisCell (D) and every form of Cellina, fitted and predicted on Cellina's own graph (top), as in its benchmark, and on ours (bottom, Sec. S3.3). The forms are as in Fig. 7, with a star marking Cellina without its domain adversary. DisCell is the same in both blocks. Values are means over three seeds with the standard deviation in parentheses, and bold marks the best mean of each block.

| Task | D | C | C′ | C″ | C* | C′* | C″* |
|---|---|---|---|---|---|---|---|
| **On Cellina's own graph** | | | | | | | |
| *Ovarian FFPE* | | | | | | | |
| Tumor | **0.23** (.03) | 0.50 (.06) | 0.51 (.06) | 0.35 (.07) | 0.48 (.01) | 0.54 (.01) | 0.32 (.02) |
| Fibroblasts | **0.37** (.05) | 0.84 (.07) | 0.82 (.07) | 0.71 (.01) | 0.88 (.14) | 0.85 (.14) | 0.70 (.10) |
| Macrophages | **0.45** (.02) | 1.06 (.13) | 1.09 (.15) | 0.80 (.06) | 1.34 (.04) | 1.33 (.04) | 1.16 (.02) |
| *Lung FFPE* | | | | | | | |
| Tumor | **0.19** (.02) | 0.46 (.02) | 0.47 (.02) | 0.46 (.02) | 0.44 (.02) | 0.44 (.02) | 0.44 (.02) |
| T/NK cells | **0.33** (.00) | 0.48 (.04) | 0.47 (.03) | 0.48 (.05) | 0.69 (.08) | 0.68 (.11) | 0.64 (.04) |
| Interst. macrophages | **0.43** (.03) | 0.65 (.10) | 0.56 (.11) | 0.52 (.09) | 0.68 (.02) | 0.60 (.02) | 0.60 (.03) |
| *Ovarian FF* | | | | | | | |
| Tumor | 0.49 (.06) | 0.46 (.05) | 0.48 (.04) | **0.43** (.06) | 0.52 (.05) | 0.54 (.06) | 0.46 (.05) |
| Ovarian stroma | **0.84** (.03) | 1.39 (.05) | 1.33 (.05) | 0.92 (.04) | 1.38 (.03) | 1.37 (.04) | 0.94 (.03) |
| *TMA core* | | | | | | | |
| Advent. fibroblasts | 0.66 (.14) | 1.25 (.12) | 1.27 (.14) | 0.80 (.10) | 1.14 (.07) | 1.18 (.08) | **0.58** (.06) |
| **On our graph** | | | | | | | |
| *Ovarian FFPE* | | | | | | | |
| Tumor | **0.23** (.03) | 0.46 (.04) | 0.40 (.01) | 0.34 (.02) | 0.47 (.07) | 0.40 (.05) | 0.31 (.02) |
| Fibroblasts | **0.37** (.05) | 0.67 (.11) | 0.78 (.12) | 0.90 (.12) | 0.66 (.03) | 0.78 (.07) | 0.94 (.07) |
| Macrophages | **0.45** (.02) | 0.56 (.04) | 0.52 (.01) | 0.46 (.05) | 1.19 (.05) | 0.99 (.04) | 0.91 (.03) |
| *Lung FFPE* | | | | | | | |
| Tumor | **0.19** (.02) | **0.19** (.03) | 0.25 (.03) | 0.30 (.03) | 0.35 (.05) | 0.38 (.01) | 0.39 (.01) |
| T/NK cells | **0.33** (.00) | 0.41 (.09) | 0.48 (.10) | 0.50 (.09) | 0.48 (.03) | 0.49 (.01) | 0.51 (.02) |
| Interst. macrophages | 0.43 (.03) | **0.39** (.06) | 0.40 (.08) | 0.42 (.09) | 0.48 (.03) | 0.48 (.04) | 0.49 (.05) |
| *Ovarian FF* | | | | | | | |
| Tumor | 0.49 (.06) | 0.40 (.03) | 0.35 (.02) | **0.30** (.02) | 0.41 (.04) | 0.39 (.02) | 0.35 (.01) |
| Ovarian stroma | 0.84 (.03) | 0.82 (.06) | 0.84 (.06) | **0.81** (.05) | 0.85 (.03) | 0.88 (.03) | 0.83 (.04) |
| *TMA core* | | | | | | | |
| Advent. fibroblasts | 0.66 (.14) | 0.98 (.12) | 0.93 (.11) | 0.82 (.12) | 1.17 (.08) | 0.74 (.06) | **0.51** (.07) |

Table S7: The ridge regression of Fig. 7, a reference that moves no cells and cannot separate spillover. It regresses expression on neighbor composition on our graph and does not depend on the seed. The scores are those of Tables S5 and S6, the correlation (higher is better) and error (lower is better) of log fold changes on the 50 most changed genes.

| Task | Correlation | Error |
|---|---|---|
| *Ovarian FFPE* | | |
| Tumor | 0.89 | 0.24 |
| Fibroblasts | 0.95 | 0.85 |
| Macrophages | 0.93 | 0.54 |
| *Lung FFPE* | | |
| Tumor | 0.78 | 0.10 |
| T/NK cells | 0.99 | 0.21 |
| Interst. macrophages | 0.99 | 0.25 |
| *Ovarian FF* | | |
| Tumor | 0.52 | 0.51 |
| Ovarian stroma | 0.92 | 0.78 |
| *TMA core* | | |
| Advent. fibroblasts | 0.97 | 0.44 |

[22] Shaked Brody, Uri Alon, and Eran Yahav. "How Attentive are Graph Attention Networks?" In: *International Conference on Learning Representations*. 2022.
